# *Collagen-binding IL-12 expressing* STEAP1 CAR-T cells reduce toxicity and eradicate mouse prostate cancer in combination with checkpoint inhibitors

**DOI:** 10.1101/2025.03.19.644145

**Authors:** Koichi Sasaki, Vipul Bhatia, Yuta Asano, Jakob Bakhtiari, Pooja Kaur, Chuyi Wang, Takumi Matsuo, Maria Nikolaidi, Po-Chuan Chiu, Olivier Dubois, Truman Chong, Gerardo Javier, Saul J. Priceman, Aude G. Chapuis, John K. Lee, Jun Ishihara

**Affiliations:** Department of Bioengineering, Imperial College London, 86 Wood Lane, London, W12 0BZ, UK; Human Biology Division, Fred Hutchinson Cancer Center, 1100 Fairview Ave N, Seattle, WA 98109, USA; Division of Hematology/Oncology, Department of Medicine, David Geffen School of Medicine at UCLA. 11-934 Factor Building, 700 Tiverton Drive, Los Angeles, CA 90095, USA; Translational Science and Therapeutics Division, Fred Hutchinson Cancer Center, 1100 Fairview Ave N, Seattle, WA 98109, USA; Department of Hematology and Hematopoietic Cell Transplantation, City of Hope, 1500 East Duarte Road, Duarte, CA 91010, USA; Department of Immuno-Oncology, Beckman Research Institute of City of Hope, 1500 East Duarte Road, Duarte, CA 91010, USA; Division of Medical Oncology, University of Washington, 1959 NE Pacific Street, Seattle, WA 98195, USA; Department of Pathology and Laboratory Medicine, University of Washington, 1959 NE Pacific Street, Seattle, WA 98195, USA; Exploratory Oncology Research & Clinical Trial Center (EPOC), National Cancer Center, Chiba, 104-0045, Japan

## Abstract

Immunosuppressive microenvironments, the lack of immune infiltration, and antigen heterogeneity pose significant challenges for chimeric antigen receptor (CAR)-T cell therapies to tackle solid tumors. CAR-T cells were armed with immunostimulatory payloads, such as interleukin-12 (IL-12), to overcome this issue, but faced intolerable toxicity during clinical development. Here, we show that collagen-binding domain-fused IL-12 (CBD-IL-12) was retained within syngeneic murine prostate tumors, after secretion from CAR-T cells targeting human six transmembrane epithelial antigen of the prostate 1 (STEAP1). This led to equivalently high intratumoral interferon-γ levels without hepatotoxicity and infiltration of T cells into non-target organs, compared with unmodified IL-12. Both innate and adaptive immune compartments were dramatically activated and recognized diverse tumor antigens after CBD-IL-12 CAR-T cell treatment. Combination immunotherapy of CBD-IL-12 CAR-T cells and immune checkpoint inhibitors eradicated large tumors in an established prostate cancer model, without pre-conditioning chemotherapy. The therapy generated anti-tumor immunological memory. CBD-fusion to potent yet toxic payloads of CAR-T therapy may remove obstacles to their clinical translation towards elimination of solid tumors.

---

Chimeric antigen receptor (CAR)-T cell therapy has demonstrated remarkable efficacy against previously incurable blood cancers^1–3^. However, the therapy has not yet been very effective against solid cancers. The immunosuppressive tumor microenvironment (TME) limits infiltration and activation of CAR-T cells and endogenous anti-tumor immune cells^4^. Furthermore, antigen heterogeneity in solid tumors poses additional difficulties for CAR-T cells, which are designed to recognize specific antigens to lyse cancer cells^4,5^.

Interleukin-12 (IL-12) is a cytokine that can remodel the immunosuppressive, “cold” TME in multiple cancer types for favorable therapeutic outcomes^6^. IL-12 is a potent inducer of interferon-γ (IFN-γ)^6,7^, which promotes trafficking and infiltration of T, NK and NKT cells into tumors through induction of chemokines, such as C-X-C motif chemokine ligand 9 (CXCL9)^8,9^. IL-12 induces proliferation of T and NK cells, and enhances their cytotoxic capacity by transcriptional reprogramming^10^. IFN-γ increases major histocompatibility complex (MHC)-I expression^11^ and matures cross-presenting dendritic cells (DCs)^10,12^ to promote cellular immunity against diverse tumor-associated antigens. IL-12 also reduces myeloid-derived suppressor cell (MDSC) numbers by facilitating their maturation to DCs and macrophages^13^.

Despite great promise, the clinical application of IL-12 has been hampered by dose-limiting immune-related adverse events (irAEs)^14,15^, strongly suggesting the need for optimized drug delivery systems. Researchers have attempted to use the ability of tumor-reactive T cells to migrate into tumor tissue for delivery of IL-12^16,17^. The nuclear factor of activated T cells (NFAT)-responsive promoter enables the engineered T cells to express payloads (i.e. anti-tumor biologics) upon recognition of tumor antigen, thereby restricting systemic concentration of the payload^18^. NFAT promoter-driven induction of unmodified IL-12 was reportedly safer than constitutive expression of IL-12^19^. However, in a Phase I trial, 8 of 16 metastatic melanoma patients who received ≥300 million NFAT-IL-12-equipped tumor-infiltrating lymphocytes exhibited grade 3 or 4 hepatotoxicity, which hampered clinical development^20^. Additional technological advances to counteract the toxicity of IL-12 are required to combine IL-12 with CAR-T cells for clinical use against solid tumors.

We previously showed that both intravenously- and intratumorally-injected A3 collagen-binding domain (CBD) of von Willebrand factor (vWF) effectively accumulates in tumors, due to tumor-specific collagen accessibility^21^. A recombinant fusion of CBD to IL-12 (CBD-IL-12) improved both safety and anti-tumor efficacy^22^. CBD-IL-12 accumulated in the tumor stroma due to high collagen expression and collagen exposed in the dysregulated tumor vasculature. In immunologically cold melanoma and breast cancer models, intravenous CBD-IL-12 induced long-lasting elevation of intratumoral IFN-γ levels, compared to unmodified IL-12. CBD fusion to IL-12 significantly reduced liver damage marker alanine transaminase (ALT) activity and serum IFN-γ.

Metastatic castration-resistant prostate cancer (mCRPC) is a largely incurable solid tumor, with a median overall survival of only three years^23^. We have recently identified a promising tumor-associated antigen, six transmembrane epithelial antigen of the prostate 1 (STEAP1), which is more broadly expressed than prostate-specific membrane antigen (PSMA) in mCRPC^24^. We have developed 2^nd^ generation CAR-T cells, using 4-1BB-based mouse and human CARs against human STEAP1 (hSTEAP1), which showed significant, yet non-curative efficacy in multiple prostate cancer models in mice^24^. Intravenously injected CBD-IL-12 protein and STEAP1 CAR-T cells synergized to extend survival of prostate cancer-bearing mice^24^.

Here, we further engineered STEAP1 CAR-T cells to express CBD-IL-12 upon STEAP1 recognition, to enhance tumor-specific immune responses. We hypothesized that fusion of CBD to IL-12 as a cargo would decrease the systemic toxicity associated with armoring T cells with unmodified IL-12.

## Results

### CBD-IL-12-armored STEAP1 CAR-T cells efficiently killed cancer cells in vitro

We designed single-chain (sc) mouse IL-12 variants with different positions of CBD-fusion (Fig. 1A, table S1), produced the constructs in HEK293F cells and purified them by affinity His-tag chromatography and size exclusion chromatography. The purity of IL-12 variants was confirmed, using sodium dodecyl sulfate-polyacrylamide gel electrophoresis (SDS-PAGE) (Fig. 1B). Signal transducer and activator of transcription 4 (STAT4) phosphorylation assays revealed that all CBD-IL-12 variants have potent yet slightly decreased EC50 values, compared to the unmodified scIL-12, presumably due to steric hinderance of the CBD fusion (Fig. 1C). We next designed gamma-retroviral vectors incorporating the mouse CAR targeting hSTEAP1^24^ and an IL-12 variant (Fig. 1D). Expression of the two genes is separately driven by a constitutive spleen focus-forming virus (SFFV) promoter and an NFAT-responsive promoter, respectively. Gamma-retroviruses were produced and used to transduce mouse primary T cells purified from spleens of male mice. Mouse primary T cells were transduced with IL-12 variant-armored CAR constructs; we noted some reduction in transduction efficiency in proportion to the length of the transgenes, whereas the conventional hSTEAP1 CAR construct transduced up to ∼80% of T cells, consistent with our previous study^24^ (Fig. 1E, S1A). A vector containing a constitutive enhanced green fluorescent protein (EGFP) expression cassette and a NFAT-driven CBD-IL-12-CBD expression cassette (CBD on both ends) was also prepared as a control to test IL-12 production triggered by CAR-antigen interaction. The transduction efficiency of EGFP + NFAT-CBD-IL-12-CBD was similar to that of hSTEAP1 CAR + NFAT-CBD-IL-12-CBD (Fig. S1B).

**Figure 1.**
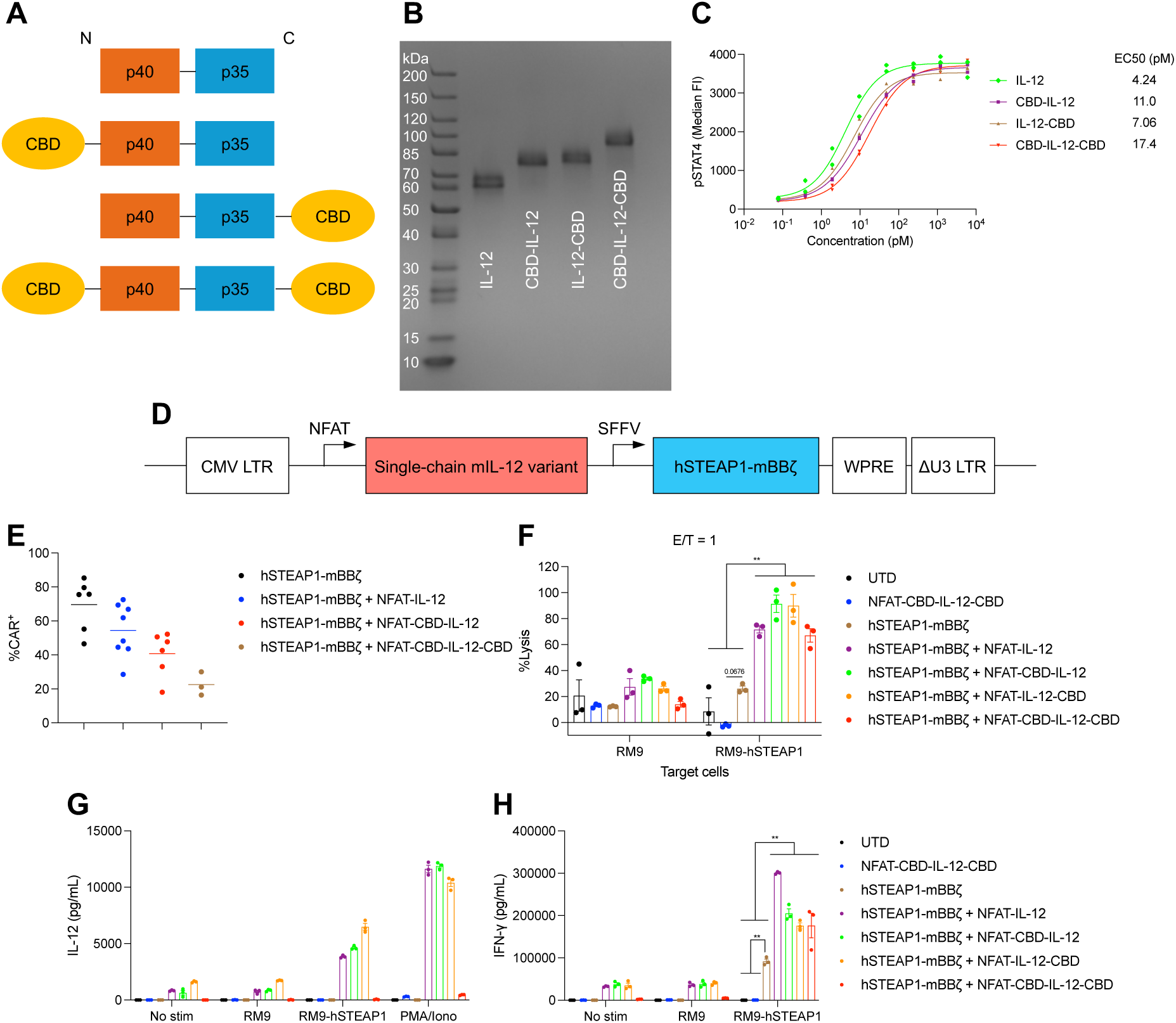
Characterizations of CAR-T cells armored with a collagen-binding IL-12 in vitro. (A) Schematic of designs and configurations of single-chain (sc) mouse IL-12 variants used in this study. (B) scIL-12 variants were analyzed by SDS-PAGE under reducing conditions with Coomassie blue staining. (C) Dose-response relationship of phosphorylated STAT4 (pY693) with scIL-12 variants in preactivated primary mouse CD8^+^ T cells (n = 2 per condition, technical duplicates). EC50, half-maximal effective concentration. (D) Schematic of self-inactivating gamma-retroviral vectors used in this study. (E) Primary mouse T cells transduced with the indicated gamma-retroviral vectors were assessed for the expression of the CAR targeting STEAP1 by flow cytometry. Individual values (biological replicates) with mean. (F) RM9-hSTEAP1 cells or unmodified RM9 cells labelled with Calcein-AM were co-cultured with CAR-T cells at the Effector/Target ratio of 1:1 for 24 h (Mean ± SEM). (G, H) Primary mouse CAR-T cells (50000 CAR-T cells/well) were left untreated (No stim), stimulated with RM9 cells, RM9-hSTEAP1 cells or phorbol 12-myristate 13-acetate and ionomycin (PMA/Iono) for 24 h. (G) Secreted IL-12 variants and (H) IFN-γ were quantified by ELISA (Mean ± SEM). Statistical analyses were performed using (F, H) one-way ANOVA with Tukey’s test (within RM9-hSTEAP1 co-culture samples). **P < 0.01.

All of the CAR-T cells lysed RM9 mouse prostate cancer cells expressing hSTEAP1 and firefly luciferase (fluc), whereas unmodified RM9 cells were not lysed (Fig. 1F, S2). CAR-T cells armored with any of the IL-12 variants showed significantly higher cytotoxicity against RM9-hSTEAP1-fluc cells than the conventional hSTEAP1 CAR-T cells, suggesting that the enhanced cytolytic capacity is due to autocrine IL-12 signaling^10^. CAR-T cells equipped with one CBD fused to IL-12 secreted the payload upon stimulation with hSTEAP1 expressed on the tumor cells, or phorbol 12-myristate 13-acetate plus NFAT-activating ionomycin (PMA/Iono) (Fig. 1G). However, the amount of secreted CBD-IL-12-CBD was significantly lower than measured for other IL-12 variants. Nonetheless, IL-12-armored CAR-T cells (including CBD-IL-12-CBD CAR-T cells) secreted significantly higher amounts of IFN-γ than unarmored CAR-T cells, consistent with IL-12-mediated autocrine stimulation of the armored CAR-T cells (Fig. 1H). These results demonstrate that the IL-12 variant-expressing hSTEAP1 CAR-T cells can lyse hSTEAP1-positive cancer cells and secrete the payload in an antigen-dependent manner.

### hSTEAP1 CAR-T cells armored with a collagen-binding IL-12 construct demonstrate anti-tumor efficacy without pre-conditioning

We tested the therapeutic efficacy of the conventional hSTEAP1 CAR-T cells against subcutaneous, syngeneic RM9-hSTEAP1-fluc prostate cancer (Fig. 2A-D). 15 million viable T cells were intravenously administered without chemotherapy pre-conditioning on day 4 after tumor inoculation (75.3% of which were the hSTEAP1 CAR-positive). Compared to untransduced (UTD) T cells, hSTEAP1 CAR-T cells significantly delayed the tumor growth (Fig. 2A, B) and extended survival (Fig. 2C, median survival of 17 days versus 13 days), without weight loss (Fig. 2D). However, complete response (CR) was not observed. Next, we treated the same tumor model to screen the collagen-binding IL-12-armored CAR constructs in vivo (Fig. 2E-I, S3). 15 million viable T cells were intravenously administered without pre-conditioning on day 4, before blood was collected from tail veins on day 7, 11 and 14 (Fig. 2E). All the armored CAR-T treatments clearly delayed tumor growth (Fig. 2F, S3) and prolonged survival by more than two-fold, compared to UTD T cells (Fig. 2G). One of five mice treated with CBD-IL-12 CAR-T cells showed CR, as did two of six mice treated with IL-12-CBD CAR-T cells, whereas tumors were not eradicated in any mice treated with CBD-IL-12-CBD CAR-T cells. Body weight loss was not observed in any of the treatment groups (Fig. 2H). There was a non-statistically significant trend towards increased serum IFN-γ levels in mice treated with CBD-IL-12- or IL-12-CBD-armored CAR-T cells on day 11, compared to mice treated with UTD T cells (Fig. 2I). Taken together, these results indicate that fusing one CBD to either the N-terminus or C-terminus of IL-12 is more therapeutically effective than two CBDs, probably due to the poor secretion of CBD-IL-12-CBD from primary mouse CAR-T cells (Fig. 1G).

**Figure 2.**
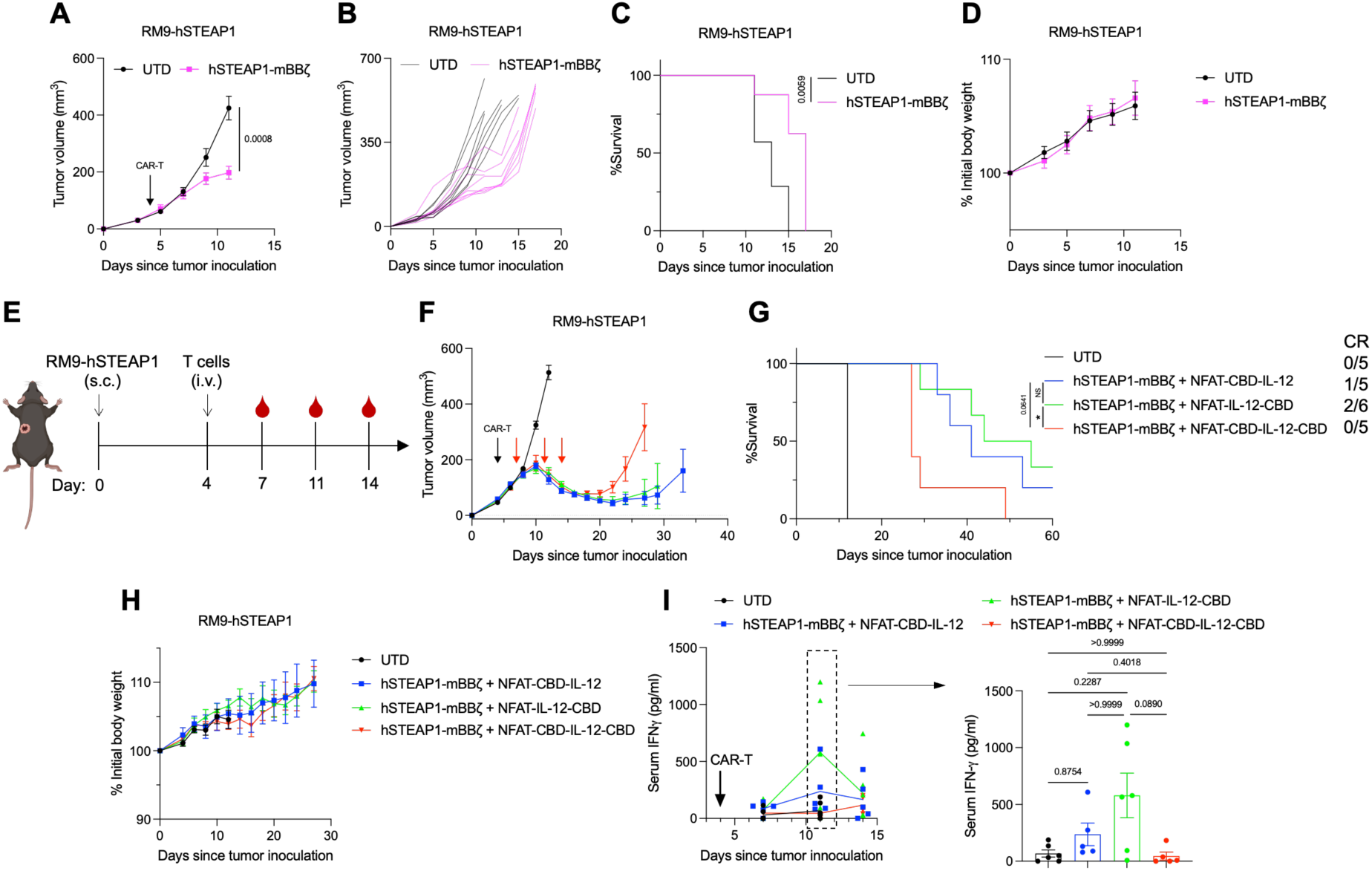
Characterizations of CAR-T cells armored with a collagen-binding IL-12 in vivo. Male C57BL6/J mice received subcutaneous injection of RM9-hSTEAP1-fluc (5×10^5^) on day 0. 15 million T cells were intravenously administered on day 4. (A-D) 75.3% CAR^+^ in hSTEAP1-mBBζ. (E-I) 54.1% CAR^+^ in hSTEAP1-mBBζ + NFAT-CBD-IL-12, 62.8% CAR^+^ in hSTEAP1-mBBζ + NFAT-IL-12-CBD and 52.3% CAR^+^ in hSTEAP1-mBBζ + NFAT-CBD-IL-12-CBD. (A, F) Average tumor volumes (mean ± SEM). (B) Individual tumor growth curves. (C, G) Survival rates. CR, complete response. (D, H) Body weight changes normalized to the body weights on day 0 (mean ± SEM). (E) Experimental schedule. (I) Blood samples were collected from tail vein on days 7, 11 and 14. Serum IFN-γ concentrations were quantified by ELISA (mean ± SEM). (A-C) UTD, n = 7; hSTEAP1-mBBζ, n = 8. (D-G) UTD, hSTEAP1-mBBζ + NFAT-CBD-IL-12 and hSTEAP1-mBBζ + NFAT-CBD-IL-12-CBD, n = 5; hSTEAP1-mBBζ + NFAT-IL-12-CBD, n = 6. Statistical analyses were performed using (A) two-tailed Welch’s t-test, (C, G) log-rank (Mantel-Cox) test or (I) Kruskal-Wallis test followed by Dunn’s multiple comparison (non-parametric data). *P < 0.05, **P < 0.01, NS not significant.

### Secreted CBD-IL-12 demonstrates enhanced localization to tumor

We selected N-term fusion CBD-IL-12-expressing CAR-T cells for detailed comparisons with IL-12-armored CAR-T cells based on characterizations of the armored CAR-T constructs in vitro and in vivo. We first quantified the IL-12 secreted from CAR-T cells in vivo. Mice bearing hSTEAP1^+^ prostate cancers (Fig. S2) were treated with CAR-T cells. Tumor, heart, lung, liver, spleen, kidney and serum samples were obtained 2-4 days after CAR-T cell administration, for quantification of IL-12 protein. Without the administration of IL-12-armored CAR-T cells, IL-12 was barely undetectable in tumor, heart, lung and spleen (Fig. S4A, B). On day 8, a significantly higher level of intratumoral IL-12 was detected in mice treated with CBD-IL-12-expressing CAR-T cells, compared to IL-12-expressing CAR-T-treated mice (Fig. 3A). In contrast, serum concentrations of IL-12 were significantly lower in CBD-IL-12 CAR-T-treated mice than in IL-12 CAR-T-treated mice on day 8 (Fig. 3B). Similarly, significantly less IL-12 was detected from the spleens and hearts of CBD-IL-12 CAR-T-treated mice on day 6 (Fig. 3 C, D). Quantification of IL-12 in kidney and liver was technically challenging regardless of tumor implantation, due to background signals with large individual differences (Fig. S4B, C). The levels of IL-12 detected from lung were comparable between the IL-12 CAR-T-treated group and CBD-IL-12 CAR-T-treated group (Fig. S4D).

**Figure 3.**
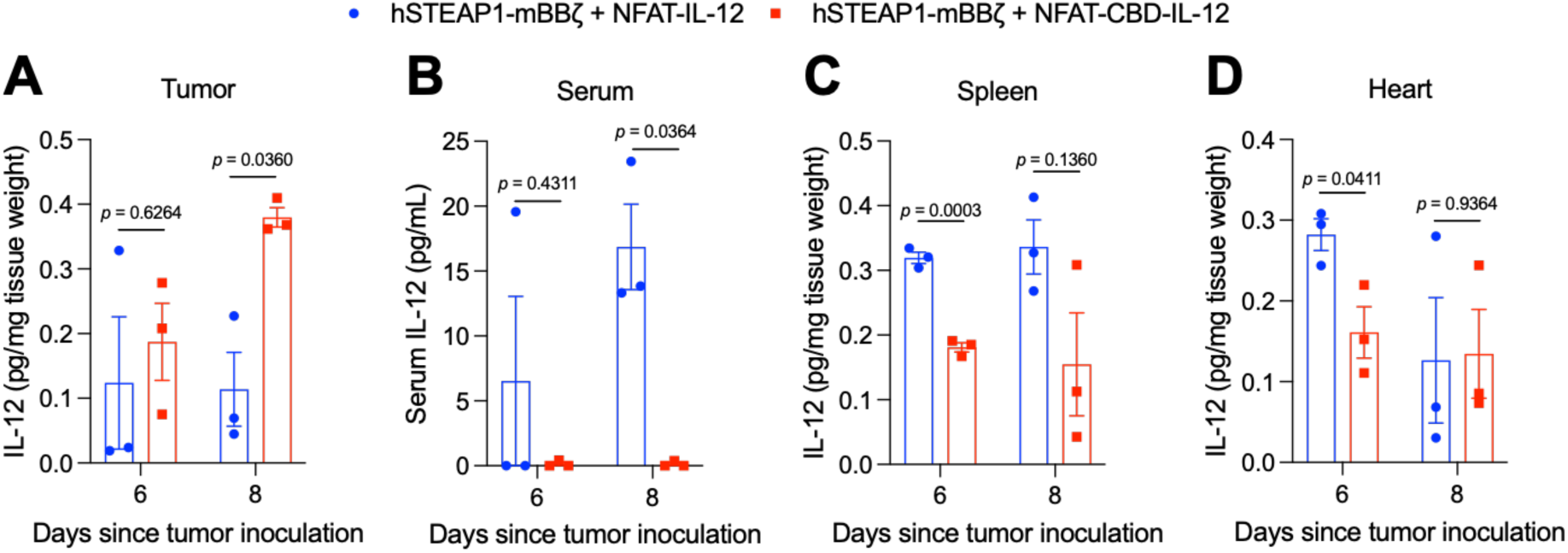
CBD fusion enhances intratumoral retention of IL-12 and reduces its systemic leakage. Male C57BL6/J mice received subcutaneous injection of RM9-hSTEAP1 (5×10^5^) on day 0. 5 million CAR^+^ T cells were intravenously administered on day 4. (A) Tumors, (B) sera and (C, D) organs were collected on indicated time-points. IL-12 was quantified by ELISA (mean ± SEM). Statistical analyses were performed using two-tailed Welch’s t-test.

### CBD-IL-12 CAR-T therapy decreases systemic toxicity

irAEs are one of the major obstacles for clinical translation of IL-12 therapies. In particular, hepatic toxicity is a common side effect in patients who receive systemic administration of recombinant human IL-12^25^. We performed blood chemistry analysis to characterize the systemic toxicity of our adoptive cell therapies in more detail. Serum concentrations of liver damage-associated ALT and alkaline phosphatase (ALP) were significantly increased after treatment with IL-12-armored CAR-T cells, but not after infusion of CBD-IL-12-armored CAR-T cells (Fig. 4A, B). IL-12-armored CAR-T cells facilitated infiltration of CD3^+^ T cells into non-target organs (kidney, lung and liver), whereas CAR-T cells and CBD-IL-12-armored CAR-T cells did not (Fig. 4 C-E). Together, these results demonstrate that the CBD/IL-12 fusion reduces systemic diffusion of IL-12, limiting the toxicity of CAR-T cells armored with this payload.

**Figure 4.**
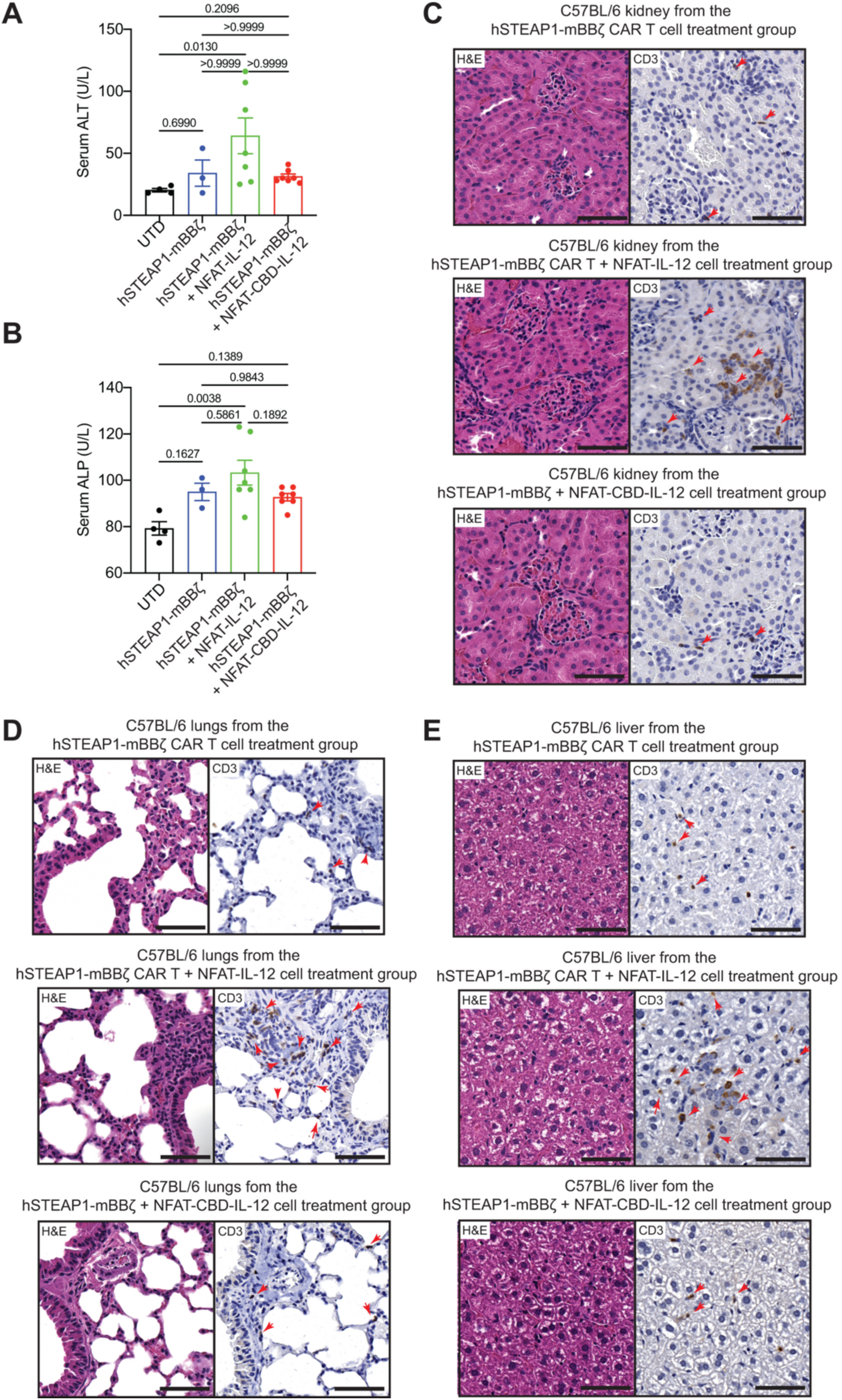
CBD decreases IL-12-related toxicity of armored CAR-T cells. Male C57BL6/J mice received subcutaneous injection of RM9-hSTEAP1 (5×10^5^) on day 0. CAR-T cells were intravenously administered on day 4. (A) Serum ALT and (B) ALP concentrations on day 12 (mean ± SEM). Histological analysis of (C) kidney, (D) lung and (E) liver on day 14. Statistical analyses were performed using (A) Kruskal-Wallis test followed by Dunn’s multiple comparison (non-parametric data) or (B) one-way ANOVA with Tukey’s test.

### CBD-IL-12 CAR-T cells inflame tumor and induce anti-tumor immune infiltrates

Tumor cell heterogeneity is the major cause of recurrence and inefficacy of CAR-T cells. Because we did not clone the RM9 cell line engineered to express hSTEAP1, there was heterogeneity in hSTEAP1 expression (Fig. S2). Indeed, hSTEAP1 CAR-T cells previously killed hSTEAP1^+^ RM9 cells in vivo but antigen-loss and recurrence occured^24^. Thus, we hypothesized that CRs seen in some of the animals (Fig. 2G) indicate that CBD-IL-12 armored CAR-T cells can induce antigen spreading. We analyzed the TME to test this hypothesis. IL-12-induced IFN-γ promotes cross-presentation by upregulating expressions of key proteins, such as MHC-I^11^. CXCL9 recruits T cells and NK cells into the TME^26^. The granulocyte macrophage colony-stimulating factor (GM-CSF) cytokine is also induced by IL-12, facilitating the maturation of cross-presenting DCs^12^ that have a major role in T cell-mediated anti-tumor immunity^27,28^. We found that IL-12- and CBD-IL-12-armored CAR-T cells upregulate intratumoral IFN-γ, CXCL9 and GM-CSF, compared with UTD and unarmored CAR-T treatments (Fig. 5A-C).

**Figure 5.**
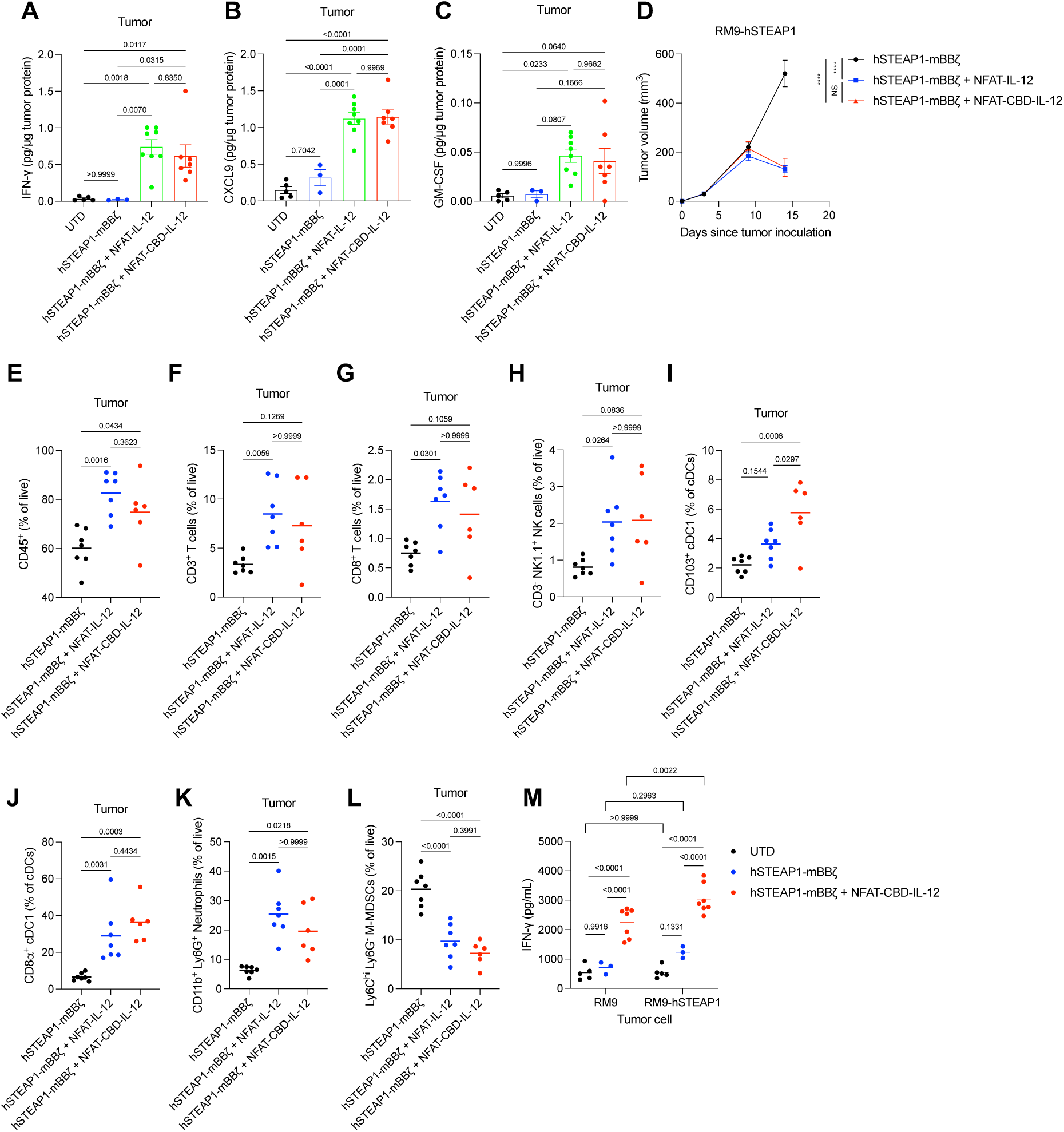
CBD-IL-12-armored CAR-T cells exert anti-tumor effect by fueling innate and adaptive immunity. Male C57BL6/J mice received subcutaneous injection of RM9-hSTEAP1 (5×10^5^) on day 0. CAR-T cells were intravenously administered on day 4. (A-C) Intratumoral concentrations of (A) IFN-γ, (B) CXCL9 and (C) GM-CSF on day 12 (mean ± SEM). (D-L) Tumors were collected on day 14 followed by flow cytometric analysis. (D) Tumor volumes (mean ± SEM). hSTEAP1-mBBζ, hSTEAP1-mBBζ + NFAT-IL-12, n = 7; hSTEAP1-mBBζ + NFAT-CBD-IL-12, n = 6. (E) Percentage of CD45^+^ cells, (F) CD3^+^ T cells, (G) CD8^+^ T cells and (H) CD3^-^ NK1.1^+^ NK cells within live cells (mean). Percentage of (I) CD103^+^ and (J) CD8α^+^ DCs among cDCs. Percentage of (K) CD11b^+^ Ly6G^+^ neutrophils and (L) Ly6C^hi^ Ly6G^-^ M-MDSCs within live cells (mean). (M) Splenic T cells were isolated and co-cultured with hSTEAP1-positive and negative RM9 prostate cancer cells. IFN-γ was quantified (mean). Statistical analyses were performed using (A-E, I, J, L) one-way ANOVA with Tukey’s test, (F-H, K) Kruskal-Wallis test followed by Dunn’s multiple comparison (non-parametric data) or (M) two-way ANOVA followed by Šídák’s multiple comparisons. ****P < 0.0001, NS not significant.

We also characterized the immune cell infiltrates in the RM9-hSTEAP1 prostate cancer model. Tumors were collected 10 days after the administration of CAR-T cells, when IL-12- and CBD-IL-12-armored CAR-T cells elicited significantly larger anti-tumor responses than unarmored CAR-T cells (Fig. 5D); intratumoral immune cells were analyzed by flow cytometry. In accordance with the observed increase of intratumoral CXCL9, CBD-IL-12 armored CAR-T therapy increased the frequency of CD45^+^ leukocytes, CD3^+^ T cells, CD8^+^ T cells and CD3^-^NK1.1^+^ NK cells, compared with unarmored CAR-T cells (Fig. 5E-H), and the increased frequencies clearly correlated with a lower tumor burden (Fig. S5A-D). CBD-IL-12 CAR-T cells increased cross-presenting conventional type 1 DCs (cDC1s), including CD103^+^ cDC1s and CD8α^+^ cDC1s within the CD11c^+^MHC-II^+^ cDC population (Fig. 5I, J), in accordance with the increase of intratumoral GM-CSF (Fig. 5C). CBD-IL-12 armored CAR-T cells significantly increased the frequency of neutrophils, which play a crucial role in eliminating antigen-negative cancer cells following adoptive transfer of antigen-specific T cells^29^ (Fig. 5K). CBD-IL-12-armored CAR-T cells significantly reduced monocytic MDSCs (M-MDSCs), an especially suppressive subset of MDSCs^30^ (Fig. 5L), and M-MDSCs frequencies positively correlated with tumor burdens (Fig. S5E). Splenic T cells derived from mice treated with CBD-IL-12-armored CAR-T cells produced a significantly higher amount of IFN-γ when co-cultured with RM9 lacking hSTEAP1 expression (Fig. 5M), suggesting T cell responses to antigens other than hSTEAP1. Collectively, CBD-IL-12-armored CAR-T cells change the composition of immune infiltrates in the tumor towards an anti-tumor state, which could explain the CRs seen in some of the treated mice.

### CBD-IL-12 CAR-T cells remodel the intratumoral transcriptional landscape to enhance anti-tumor immunity

Limited trafficking and infiltration of CAR-T cells in solid tumors and immunosuppressive TME are major challenges to the efficacy of current therapies. We used NanoString GeoMx Digital Spatial Profiler to perform spatial transcriptomics analysis and assess whether the entire TME can be remodeled by CBD-IL-12 CAR-T cells. We selected 12 regions of interest (ROIs) across three (unarmored CAR-T cells) or four (CBD-IL-12-armored CAR-T cells) different tumors. We observed infiltration of CD45^+^ immune cells throughout the tissues and reduction of tumor regions (pan-cytokeratin^+^) in CBD-IL-12 CAR-T-treated samples (Fig. 6A, B). Immunohistochemistry (IHC) staining confirmed enhanced infiltration of CD3^+^ T cells in CBD-IL-12 CAR-T-treated tumors. Principal component analysis (PCA) showed that tumor tissues treated with one of the two different therapies displayed markedly different characteristics except ROI-S1-12 (Fig. 6C). Gene set enrichment analysis (GSEA) revealed that CBD-IL-12-armored CAR-T cells activated the IL-12 pathway, and activated antigen processing and presentation by MHC-I (Fig. 6D, E). Enrichment of genes in the IL-12 pathway was not statistically significant (p = 0.059), which could be explained by the fact that only 17 genes in the dataset were annotated to this pathway. Indeed, Fisher’s exact test showed statistical significance of IL-12 pathway enrichment in differentially expressed genes (p = 0.01241). Collectively, the data suggest that CBD-IL-12-armored CAR-T cells facilitate immune cell infiltration and activation throughout the tumor, to overcome the cold TME.

**Figure 6:**
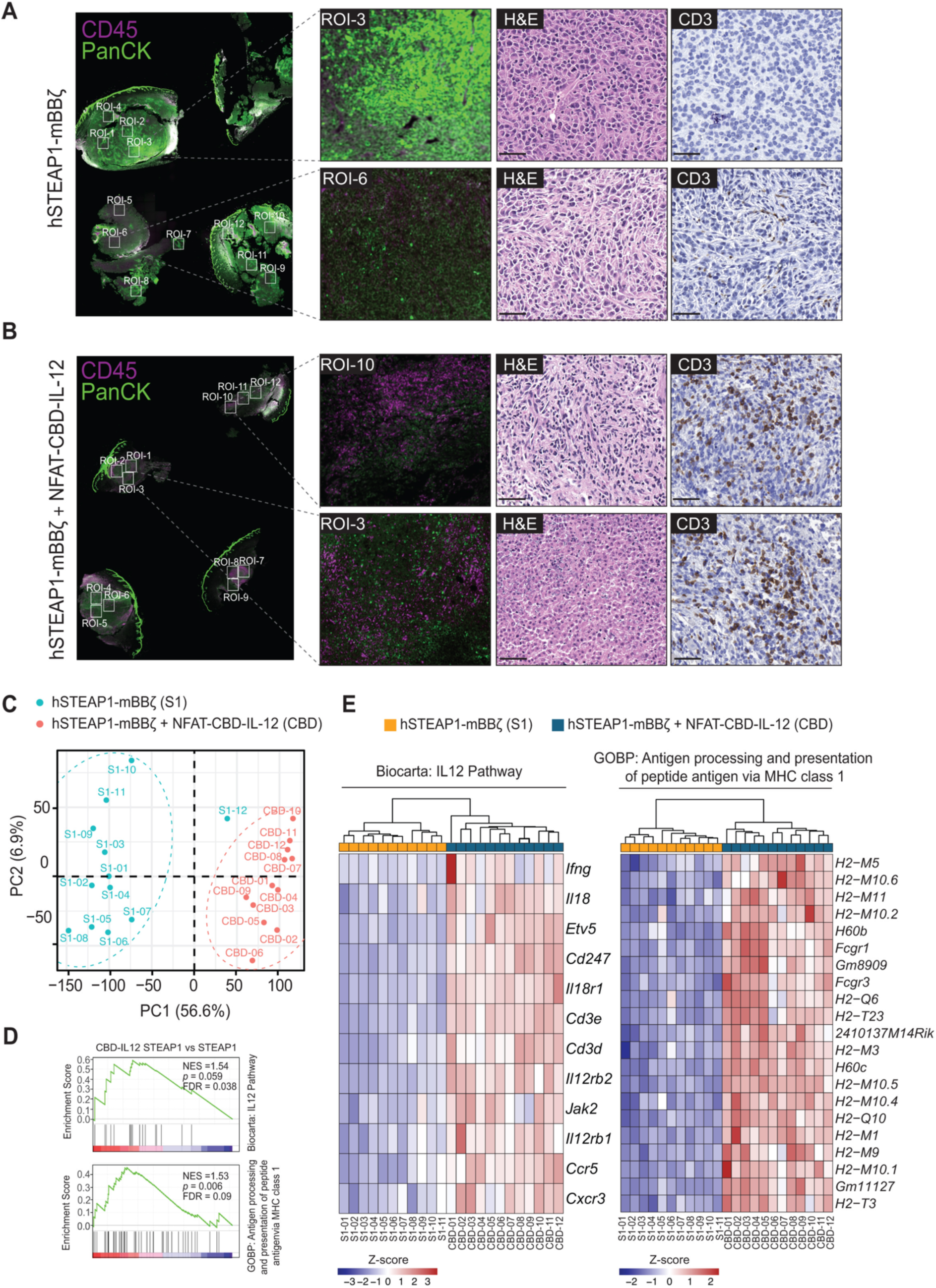
Spatial Transcriptome analysis shows CBD-IL-12 mediated changes in tumor microenvironment. Region of interest from (**A**) hSTEAP1-mBBζ (n=3) and (**B**) hSTEAP1-mBBζ + NFAT-CBD-IL-12 (n=4) treated mice. Immunofluorescence image depicting pan-cytokeratin (panCK, green) and CD45 (magenta) stain. Representative images of sections stained with H&E and CD3 (right). (**C**) PCA plot showing distribution of selected ROIs color-coded based on treatment groups. (**D**) Gene set enrichment analysis showing enriched IL-12 pathway and antigen processing and presentation via MHC-class 1. (**E**) Heatmaps showing changes in genes involved in pathways as in (D).

### CBD-IL-12 CAR-T cells combined with immune checkpoint inhibitors (CPI) eradicate large established RM9-hSTEAP1 tumors

Since CBD-IL-12 CAR-T cells alone did not induce tumor-free CRs in the majority of mice (Fig. 2G) and tumor-infiltrating T/NK cells had elevated PD-1 and CTLA-4 expression (Fig. S6), we examined the anti-tumor efficacy of CBD-IL-12 CAR-T cells + anti-PD-1 and anti-CTLA-4 CPI combination therapy. When the therapeutic intervention was started on day 4 after tumor inoculation (tumor volumes were ∼60 mm^3^), armored CAR-T cells + CPIs achieved a 100% CR rate (Fig. S7A-C), whereas tumors eventually progressed in a fraction of mice in other treatment groups. Even when the initial CAR-T treatment was delayed until day 6 after tumor inoculation (tumor volumes were ∼120 mm^3^), CBD-IL-12 CAR-T cells + CPIs therapy cured 80% of RM9-hSTEAP1-bearing mice and significantly extended the survival of the animals, compared with standard hSTEAP1-targeting CAR-T cells + CPIs (Fig. 7A, B). These data show very strong anti-tumor efficacy of combination immunotherapy against large established tumors. All the tumor-free survivors in the study rejected subcutaneously re-challenged RM9-hSTEAP1 cells (Fig. 7C), suggesting that strong anti-tumor immune memory was established in the animals. Regarding side effects evaluation, none of the treatments caused body weight loss in the animals (Fig. 7D, S7E). Previous study using IFN-γ receptor-knockout mice indicated that IL-12-induced IFN-γ plays a central role in irAEs^31^. In this study, significantly increased serum IFN-γ was observed on day 10 in mice treated with IL-12 CAR-T + CPIs, but not in mice treated with CBD-IL-12 CAR-T + CPIs (Fig. 7E), consistent with the toxicity evaluation of the CAR-T monotherapies (Fig. 4).

**Figure 7.**
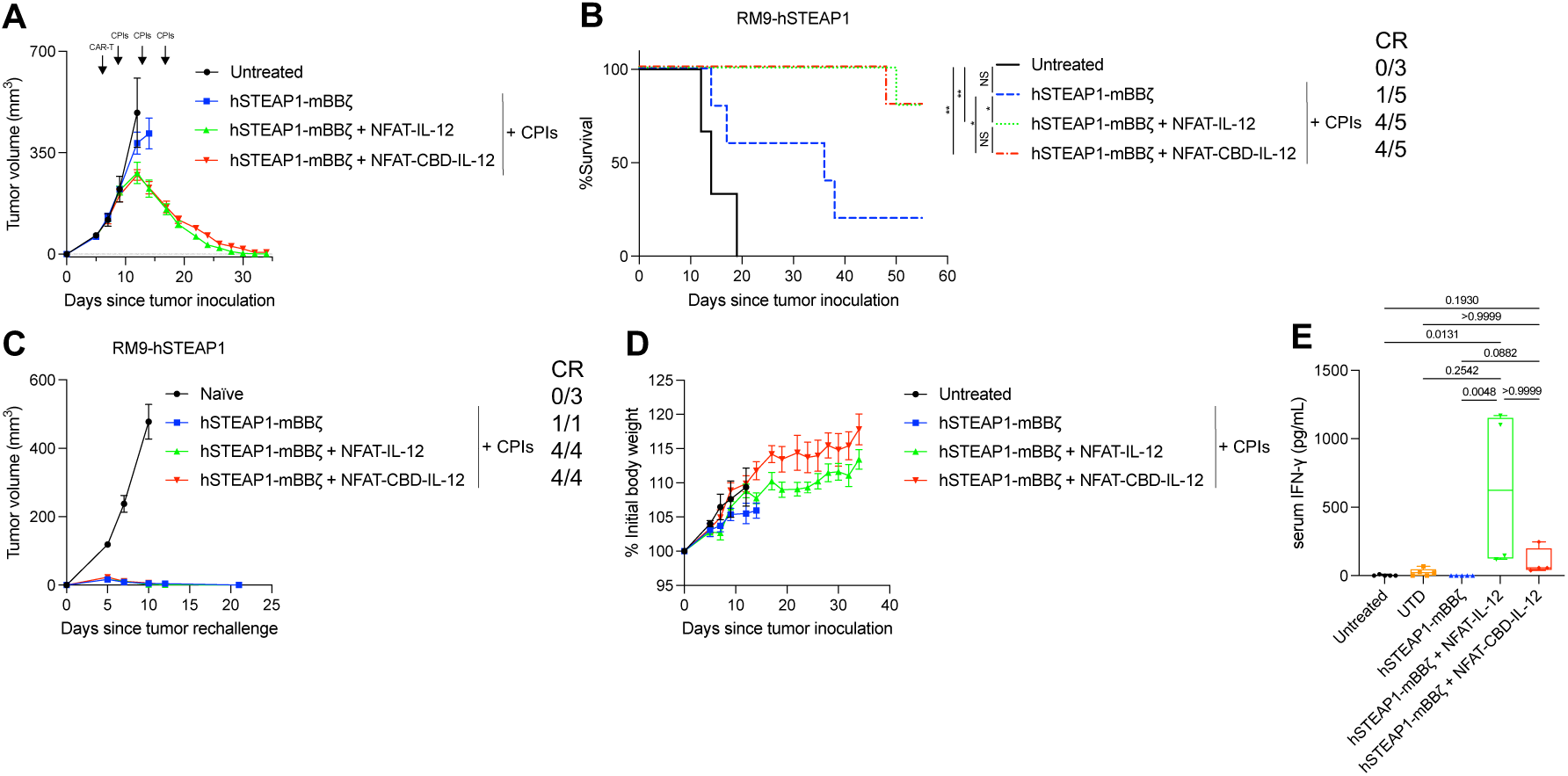
CBD-IL-12-armored CAR-T cells eradicate established RM9-hSTEAP1 tumor in combination with anti-PD-1 and anti-CTLA-4 checkpoint inhibitors. 5 million CAR^+^ T cells were intravenously administered on (A-D) day 6 or (E) day 4. Anti-PD-1 and anti-CTLA-4 antibodies were administered intraperitoneally 3 times starting at 3 days after the CAR-T administration with 4 days interval. (A) Tumor volumes were measured (Untreated, n = 3; hSTEAP1-mBBζ, hSTEAP1-mBBζ + NFAT-IL-12 and hSTEAP1-mBBζ + NFAT-CBD-IL-12, n = 5. mean ± SEM). (B) Survival rates. CR, complete response. (C) Complete responders were subcutaneously re-challenged with RM9-hSTEAP1 cells (5×10^5^) (mean ± SEM). (D) Body weight changes normalized to the body weights on day 0 (mean ± SEM). (E) Serum IFN-γ concentrations on day 10. The boxes extend from 25^th^ to 75^th^ percentiles, the center lines show median values and the whiskers extend to the minimums and maximums. Statistical analyses were performed using (B) log-rank (Mantel-Cox) test or (E) Kruskal-Wallis test followed by Dunn’s multiple comparison (non-parametric data). *P < 0.05, ** P < 0.01, NS not significant.

### Fully human CBD-IL-12 CAR-T cells kill human prostate cancer cells more efficiently than conventional CAR-T cells

After confirming that unmodified sc- and CBD-human IL-12 proteins could be transiently expressed in HEK293F cells (Fig. S8), we generated lentiviral vectors for arming human T cells, by replacing the murine CAR and CBD-IL-12 sequences with their human counterparts. We transduced Jurkat cells with the vectors to express a hSTEAP1 CAR bearing human CD3ζ and human 4-1BB signaling domains, and NFAT-driven CBD-human IL-12 (Fig. S9A). The transduced Jurkat cells secreted an IL-12 variant upon PMA/Iono stimulation (Fig. S9B). Furthermore, we successfully transduced primary human T cells derived from healthy-donor PBMC with our vectors (Fig. 8A). Human CBD- or unmodified IL-12 CAR-T cells produced IL-12 protein upon co-culture with a hSTEAP1^+^ human prostate carcinoma cell line, 22Rv1, but not when they were co-cultured with 22Rv1 hSTEAP1 KO (Fig. 8B). These IL-12 armored CAR-T cells showed better in vitro killing against 22Rv1, compared with those transduced with CAR alone (Figure 8C). The CAR-T cells did not kill 22Rv1 hSTEAP1-KO cells, demonstrating that our system is highly specific to hSTEAP1. These results indicate that the CBD-IL-12-expression platform technology is applicable to human T cells.

**Figure 8.**
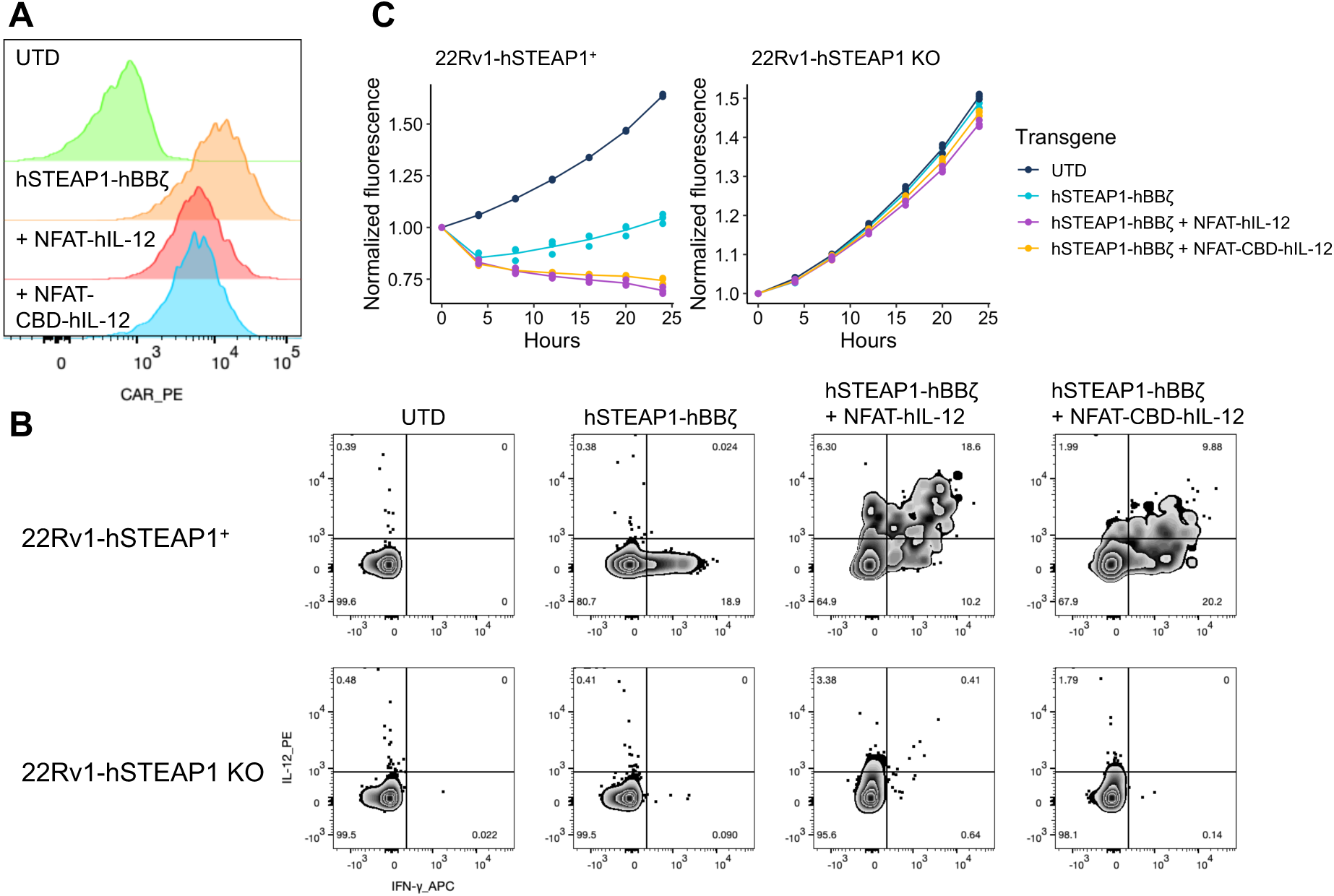
Primary human CBD-IL-12 CAR-T cells efficiently kill human prostate cancer cells and express CBD-IL-12. (A) CAR expression on expanded primary human T cells was detected on the same day of the functional assays by flow cytometry. (B) Cytokine production upon overnight co-culture with target cells were detected with intracellular flow staining. Representative zebra plots (a hybrid of contour and density plot) from three technical replicates are shown. (C) In vitro killing activity of human CD8 T cells transduced with indicated transgenes against hSTEAP1-positive or negative human prostate cancer target cells. Values from three technical replicate (dots) and mean values (lines) are shown.

## Discussion

The ultimate goal of cancer immunotherapy is to achieve safe and long-term effectiveness, as certain CAR-T cells are showing remarkable clinical responses against some hematological malignancies. The next milestone for CAR-T therapies is to overcome solid tumor unresponsiveness and/or tumor recurrence. We have previously reported that the 2^nd^-generation STEAP1 CAR-T cells induce tumor regression, but also face recurrence due to STEAP1 antigen loss that might be overcome by combining with a potent cytokine, such as IL-12, to efficiently activate the host immune system. We previously reported that conjugation of vWF-derived CBD to CPI antibodies (PD-L1, CTLA4), cytokines (IL-2 and IL-12), chemokine CCL4 or serum albumin improves their localization to the tumor stroma, enhancing therapeutic efficacy and decreasing side effects^21,22,32,33^. In our hands, CBD-IL-12 showed the strongest anti-tumor effects in multiple mouse models, including complete remission and immunological memory generation.

To our knowledge, this is the first time that a CAR-T payload has been used to target extracellular matrix. CBD-IL12 expression from our novel CAR-T cell therapy improved the potency of CAR-T cells against mouse prostate cancer. Compared with unmodified IL-12, the CBD fusion significantly enhanced intratumoral retention of IL-12 from the CAR-T payload and reduced systemic IL-12 exposure, also producing higher intratumoral IFN-γ levels and anti-tumor efficacy while reducing hepatotoxicity, T cell infiltration into non-target organs, and systemic IFN-γ. As side effects of IL-12-expressing adoptive T cell therapy is the major reason for clinical trial failures, our data demonstrate a crucial improvement. The tumor-homing capacity of CBD-IL-12^22^ may contribute to toxicity reduction even when CBD-IL-12 is secreted from CAR-T cells in healthy tissues, due to the leakiness of the NFAT-responsive promoter and/or low-level STEAP1 expression in irrelevant tissues. Our data demonstrate that CBD fusion to IL-12 enhances the safety of IL-12 as a cargo of adoptive T cell therapies, and CBD-based drug delivery systems could be applied to other cell therapies.

Chemotherapy pre-conditioning is widely used in combination with CAR-T therapy, to remove pre-existing immune cells and enable CAR-T cells to infiltrate and remodel the TME^34^. While effective, chemotherapy often causes hematological and systemic toxicities and increased risks of too-often lethal infections^35^. To our surprise, CBD-IL-12 CAR-T cells + CPI antibodies eradicated RM9 tumors without chemotherapy pre-conditioning, which might contribute to improving patients’ quality of life if translated to the clinic.

One of the greatest challenges in the treatment of solid malignancies is antigen heterogeneity. IL-12 is exactly the cytokine for tackling this issue by enhancing the capacity of cellular immunity to fight diverse antigens. Our flow cytometric, transcriptomic and splenic T cell restimulation data suggest that CBD-IL-12 CAR-T cells can induce antigen spreading to fight antigen heterogeneity in solid tumors.

Other approaches to reduce toxicity of IL-12 as a payload include tethering IL-12 to the CAR-T cell surface by fusing a transmembrane domain to IL-12, as we and others previously reported^36–38^. Other approaches include cell-surface tethering through fusing an antibody fragment targeting cell surface receptors^39^ and conjugation of IL-12-loaded nanoparticles onto azide-labeled CAR-T cells through click chemistry^40^. Although these methods can be powerful in reducing toxicity, they might reduce the chance for IL-12 to act in trans via directly binding to IL-12 receptor on endogenous immune cells. It will be important to see how these methodologies may differ from secreted IL-12 in terms of levels of therapeutic efficacy and mechanisms of action.

Our matrix-binding technology has several additional advantages for clinical applications to a wide range of CAR-T cell therapies against solid tumors. First, collagen is abundantly expressed across patients with multiple tumor types, likely making this technology useful for a wide variety of patients. Second, identifying a promising target antigen for each solid tumor type is a major challenge in the field^4,5^, while our versatile, collagen-targeting approach enables a payload to avoid competition with CARs. Third, CBD is unlikely to confer immunogenicity to CAR-T cells because it is of fully human origin.

In summary, this study demonstrates a protein-engineering approach to make IL-12 safer for use with CAR-T therapy. The utility of CBD-IL-12 lies in its capacity to overcome the immunosuppressive TME and antigen heterogeneity while avoiding irAEs and competition with CARs for tumor targeting, providing a high potential for clinical translation.

## Methods

### Cell lines

HEK293T (CRL-3216), RM9 (CRL-3312), MyC-CaP (CRL-3255), 22Rv1 (CRL-2505) and Jurkat (TIB-152) cells were obtained from ATCC. HEK293T, RM9 and MyC-CaP cells were maintained in DMEM medium (Gibco) supplemented with 10% FBS (Gibco), 2 mM GlutaMAX (Gibco) and 1% penicillin/streptomycin (Gibco). 22Rv1 and Jurkat cells were maintained in RPMI1640 (Gibco) supplemented with 10% FBS (Gibco), 2 mM GlutaMAX (Gibco) and 1% penicillin/streptomycin (Gibco). PLAT-E (RV-101) was obtained from Cell Biolabs and maintained in DMEM medium (Gibco) supplemented with 10% FBS (Gibco), 2 mM GlutaMAX (Gibco), 1% penicillin/streptomycin (Gibco), 1 µg/mL puromycin (Merck) and 10 µg/mL blasticidin (Stratech). RM9-hSTEAP1-fluc, 22Rv1 hSTEAP1 KO and 22Rv1 hSTEAP1 KO + rescue cells were generated in our previous work^24^. Myc-CaP-hSTEAP1-CGW cells were generated using FU-hSTEAP1-CGW vector^24^.

### Validation of hSTEAP1 expression on prostate cancer cells

Cells were resuspended in cold PBS (-) and stained with BD Horizon Fixable Viability Stain 510 (BD). After a wash with PBS (-) supplemented with 2% FBS, the cells were stained with Vandortuzumab (Invitrogen) followed by Alexa Fluor 594 AffiniPure F(ab’)_2_ Fragment Donkey anti-human IgG (H+L) (Jackson ImmunoResearch, 709-586-149). Cells were acquired using a BD FACSymphony A3 flow cytometer and data were analyzed using FlowJo (BD).

### Mice

6 to 9 weeks old male C57BL/6J mice and FVB/N mice were obtained from Charles River UK and housed at the Hammersmith Central Biomedical Services facility of Imperial College London. All the animals were handled in accordance with the 1986 Animal Scientific Procedures Act and under a United Kingdom Government Home Office-approved project license and overseen by ethical committees of Imperial College London.

### Production and purification of recombinant single-chain (sc) IL-12 variants

Optimized sequences encoding p35 and p40 connected with a glycine-serine linker were synthesized and subcloned into pcDNA3.1(+) vector by GenScript. The collagen-binding domain (A3 domain of human VWF)^41^ was fused to N-terminus, C-terminus or both of the protein to create CBD-IL-12 variants. His-tag was added to the C-terminus after the CBD. All the IL-12 variants were also subcloned into pcDNA3.1(+) vector. Recombinant IL-12 variants were produced by transient expression in HEK293F cells and purified as described previously^22^. Purity of the proteins were evaluated using SDS-PAGE as described previously^22^. Protein concentration was quantified by measuring absorbance at 280 nm using a NanoDrop One (Thermo Fisher Scientific). Full amino acid sequences of the IL-12 variants are available in the supplementary information.

### STAT4 phosphorylation assay

Primary mouse T cells were treated with IL-12 variants to evaluate phosphorylation of STAT4 as described previously^22^. In brief, CD3^+^ T cells were purified from spleens of C57BL/6J mice using MagniSort Mouse T cell Enrichment Kit (Invitrogen) according to the manufacturer’s instructions. Purified T cells were activated using T cell Activation/Expansion kit, mouse (Miltenyi Biotec) according to the manufacturer’s instructions and cultured in RPMI1640 medium (Gibco) supplemented with 10% FBS (Gibco), 2 mM GlutaMAX (Gibco), 1% MEM Non-essential amino acids (Gibco), 1 mM sodium pyruvate (Gibco) 50 µM 2-mercaptoethanol (Gibco), 1% penicillin/streptomycin (Gibco) and 10 ng/mL recombinant human IL-2 (Peprotech) for 3 days. T cell activation/expansion beads were magnetically removed, and the T cells were rested in fresh culture medium (without IL-2) overnight. T cells were seeded into 96-well V-bottom plates at 1 × 10^5^ cells/well. T cells were stimulated with the indicated concentrations of IL-12 variants at 37°C for 15 min. Cells were fixed with BD Phosflow Lyse/Fix buffer and permeabilized with BD Phosflow Perm Buffer III according to manufacturer’s instructions. Cells were stained with anti-pSTAT4 Alexa Fluor 647 (pY693, BD) and Anti-mouse CD8α BV510 (53-6.7, Biolegend) and acquired using a BD FACSymphony A3 flow cytometer. Data were analyzed using FlowJo (BD).

### CAR expression plasmids

pSIRV-NFAT-eGFP was a gift from Peter Steinberger (Addgene plasmid #118031). pHR_SFFV was a gift from Wendell Lim (Addgene plasmid #79121). hSTEAP1-BBζ CAR sequences were prepared as described previously^24^. SFFV promoter, hSTEAP1-mBBζ CAR, mouse scIL-12 variant and WPRE were subcloned into the pSIRV-NFAT-eGFP vector via in-fusion cloning (Takara Bio) as illustrated in Fig. 1D. eGFP sequence was replaced with IL-12 sequence during the process. MSCV promoter, NFAT-responsive promoter, human scIL-12 variant, WPRE and hSTEAP1-hBBζ CAR sequences were cloned into pRRLSIN vector backbone^42^.

### Production of gamma-retroviral vectors

For mouse CAR gamma-retrovirus production, PLAT-E cells (Cell Biolabs) were maintained in DMEM medium (Gibco) supplemented with 10% FBS (Gibco), 2 mM GlutaMAX (Gibco), 1% penicillin/streptomycin (Gibco), 1 µg/mL puromycin (Merck) and 10 µg/mL blasticidin (Stratech). A day prior to transfection, PLAT-E cells were resuspended in antibiotic-free medium with 10%FBS and GlutaMAX, seeded and cultured for 24 h. PLAT-E cells were transfected with a retroviral transfer vector using GeneJuice (Merck Millipore). Supernatant was harvested at 48 h and 72 h after transfection and passed through a 0.45 µm PES filter (Merck). Supernatant collected at 48 h was stored at 4°C overnight and pooled together with supernatant collected at 72 h. Pooled supernatant was concentrated by high-speed centrifugation (24000 g at 4°C for 2 h) using Avanti JXN-30 (Beckman Coulter) with JS-24.15 rotor (Beckman Coulter) and resuspended in DMEM medium (Gibco) supplemented with 10% FBS. Concentrated viral vector was immediately added to a non-treated cell culture plate pre-coated with 20 µg/mL Retronectin (Takara Bio) and centrifuged (2000 g at 32°C for 1.5 h) before transduction of T cells.

### Producion of lentiviral vectors

pMD2.G (Addgene plasmid #12259), pMDLg/pRRE (Addgene plasmid #12251) and pRSV-Rev (Addgene plasmid #12253) were gifts from Didier Trono. HEK293T cells (ATCC) maintained in DMEM medium (Gibco) supplemented with 10% FBS (Gibco), 2 mM GlutaMAX (Gibco) and 1% penicillin/streptomycin (Gibco) were used. HEK293T cells were transfected with a lentiviral transfer vector, pMD2.G, pMDLg/pRRE and pRSV-Rev.

### Manufacturing of primary mouse CAR-T cells

Splenocytes were harvested from spleens obtained from male C57BL/6J or FVB/N mice (Charles River UK). A spleen was put on a 70 µm cell strainer pre-wet with DMEM supplemented with 2% FBS and mashed using a plunger of 2.5 mL syringe (Terumo). The strainer was washed with the DMEM medium twice and centrifuged at 300 g for 5 min. After removing the supernatant, red blood cells were lysed using ACK lysing buffer (Gibco) at room temperature for 5 min. Cells were washed with excess PBS (-) and resuspended in PBS (-) supplemented with 2% FBS and 2 mM EDTA. Mouse T cells were purified using MagniSort Mouse T cell Enrichment Kit (Invitrogen) according to the manufacturer’s instructions. Purified T cells were activated using T cell Activation/Expansion kit, mouse (Miltenyi Biotec) according to the manufacturer’s instructions and cultured in RPMI1640 medium (Gibco) supplemented with 10% FBS (Gibco), 2 mM GlutaMAX (Gibco), 1% MEM Non-essential amino acids (Gibco), 1 mM sodium pyruvate (Gibco) 50 µM 2-mercaptoethanol (Gibco), 1% penicillin/streptomycin (Gibco) and 10 ng/mL recombinant human IL-2 (Peprotech). 24 h later, the activation/expansion beads were magnetically removed and 1 µg/mL anti-mouse IL-12 p40 (C17.8, Bio X Cell) was added for IL-12 armored CAR-T cells. T cells were seeded into a retronectin-treated viral vector-coated plate and centrifuged (300 g at 32°C for 10 min) and cultured for 48 h. After the 48 h of culture, the cytokine supplementation was changed from recombinant human IL-2 to 10 ng/mL recombinant human IL-7 (Peprotech) and 10 ng/mL recombinant human IL-15 (Peprotech). %CAR^+^ was assessed using either a combination of biotinylated recombinant protein L (Thermo Scientific) and streptavidin-Alexa Fluor 647 conjugate (Invitrogen) (for Fig. 2 and S3) or a combination of Biotin-SP-conjugated AffiniPure F(ab’)_2_ Fragment Goat Anti-Human IgG, F(ab’)_2_ Fragment Specific (Jackson ImmunoResearch 109-066-006) and PE Streptavidin (Biolegend) (all the other figures) on day 4. Cells were acquired using a BD FACSymphony A3 flow cytometer and data were analyzed using FlowJo (BD). CAR-T cells were resuspended in cold PBS (-), stained with Acridine Orange/Propidium Iodide (DeNovix) and counted using a CellDrop automated cell counter (DeNovix) for adjusting the concentration before use in animal experiments. CAR-T cells were intravenously injected to mice on day 5 since viral transduction.

### CAR-T cell-mediated cytotoxicity assay

1 × 10^5^ CAR-T cells/mL in the culture medium without supplementation of any interleukins were seeded into a 96-well V-bottom plate. Target cells were labelled with 1 µM Calcein-AM as described previously^43^. 1 × 10^5^ labelled target cells/mL in the culture medium with 5 mM probenecid were mixed with the CAR-T cells at 10000 target cells/well. The cells were centrifuged (300 g for 5 min at 37°C) and incubated for 24 h at 37°C in the presence of 5% CO_2_. The plate was centrifuged and 100 µL/well of the supernatant was transferred to a 96-well black F-bottom plate (Greiner Bio-One). Fluorescence intensity (Ex 490 nm, Em 530 nm) was measured using CLARIOstar Plus microplate reader (BMG Labtech). After subtracting autofluorescence derived from medium, %lysis was calculated using the following formula:

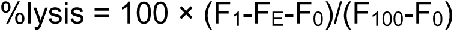

F_1_: Fluorescence intensity of the mixture of CAR-T cells and labelled target cells

F_E_: Fluorescence intensity of CAR-T cells

F_0_: Fluorescence intensity of labelled target cells

F_100_: Fluorescence intensity of labelled target cells lysed with 1% Triton X-100

### Enzyme-linked immunosorbent assay (ELISA) for detection of IL-12 and IFN-γ secreted from CAR-T cells

5 × 10^5^ CAR-T cells/mL in the culture medium (without supplementation of any interleukins) and 5 × 10^5^ target cells/mL in the same medium were seeded into a 96-well V-bottom plate (50000 CAR-T cells/well, CAR-T/Target ratio = 1), centrifuged (300 g for 5 min at 37°C) and incubated for 24 h at 37°C in the presence of 5% CO_2_. The plate was centrifuged again, and the culture supernatant was collected for quantification of IL-12 and IFN-γ using UTD kits for the mouse proteins (Invitrogen). eBioscience Cell Stimulation Cocktail (Invitrogen) was used as a cocktail of PMA and Ionomycin. For quantification of scIL-12 variants secreted from CAR-T cells, the IL-12 variants with his-tag were produced using HEK293F cells as described above and used as a standard.

### Anti-tumor efficacy of CAR-T cells against RM9-hSTEAP1

A total of 5 × 10^5^ RM9-hSTEAP1-fluc cells resuspended in 50 µL of PBS (-) were subcutaneously injected on the left side of the back of each male C57BL/6J mouse on day 0. Primary mouse CAR-T cells were intravenously injected on day 4 or day 6. The dose of CAR-T cells in each study is described in figure legends. In some studies, anti-CTLA-4 (9H10, Bio X cell) and anti-PD-1 (RMP1-14, Bio X cell) were intraperitoneally injected (100 µg each) 3 times starting at 3 days after the CAR-T administration with 4 days interval. Tumors were measured using a digital caliper and the volumes were calculated as ellipsoids (V = 4/3 × 3.14 × depth/2 × width/2 × height/2). Blood samplings from tail veins were performed at the indicated time points to assess serum concentration of IFN-γ by ELISA (Invitrogen). Mice were euthanized when tumor volume had exceeded 500 mm^3^ or tumor ulceration of more than 5 mm in diameter had been observed.

### Quantification of IL-12 in serum, tumor and major organs

RM9-hSTEAP1-fluc tumor-bearing mice received 5 million CAR-T cells on day 4 since tumor inoculation. Blood samples, tumors and major organs were collected at the indicated time points. Blood samples were incubated at room temperature for at least 1 h and centrifuged at 1000 g for 20 min at 4°C. Serum was collected in 1.5 mL protein lobind tubes (Eppendorf) and stored at -80°C until use. Parts of the removed tumors and organs were immediately put into pre-weighed Lysing Matrix D tube (MP Biomedicals) containing 1 mL of T-PER Tissue Protein Extraction Reagent (Thermo Scientific) supplemented with cOmplete EDTA-free protease inhibitor cocktail. Tubes with tissue samples were weighed again and cut into small pieces using surgical scissors. The samples were lysed using FastPrep-24 5G (MP Biomedicals) and stored at -80°C until use. IL-12 was quantified using mouse IL-12 p70 ELISA kit (Invitrogen) and normalized by total tissue weight. Recombinant CBD-mouse scIL-12 and mouse scIL-12 (Table S1) were used as standards for each protein in ELISA.

A total of 2 × 10^6^ MyC-CaP-hSTEAP1-CGW cells resuspended in 50 µL of 50% PBS (-) and 50% Phenol Red-free Matrigel (Corning) were subcutaneously injected on the left side of the back of each male FVB/N mouse. FVB/N male mice MyC-CaP-hSTEAP1-CGW tumor-bearing mice received 5 million CAR-T cells when average tumor volume was about 230 mm^3^. Tumors were collected 4 days after CAR-T cell administration and 3 fragments were obtained from each tumor tissue for the experiment. Intratumoral IL-12 was quantified as described above and normalized by total protein content. Total protein content was quantified using the Pierce BCA Protein Assay Kit (Thermo Fisher Scientific). Bovine serum albumin was used as standards.

### Systemic toxicity of armored CAR-T cell therapy

RM9-hSTEAP1-fluc tumor-bearing mice received 10 million CAR-T cells on day 4 since tumor inoculation. Terminal blood sampling by cardiac puncture for blood chemistry analysis was performed on day 12. Serum ALT and ALP were quantified using Skyla VB1 veterinary chemistry analyzer. Skyla preanesthetic panel discs were used according to the manufacturer’s instructions.

### Quantification of cytokines in CAR-T treated tumors

RM9-hSTEAP1-fluc tumor-bearing mice received 10 million CAR-T cells on day 4 since tumor inoculation. Tumors were harvested on day 12 and put into Lysing Matrix D tube (MP Biomedicals) containing T-PER Tissue Protein Extraction Reagent (Thermo Scientific) supplemented with cOmplete EDTA-free protease inhibitor cocktail. Tumors were cut into small pieces using surgical scissors and lysed using FastPrep-24 5G (MP Biomedicals). All the samples were stored at -80°C until use. Intratumoral cytokines were quantified using a LEGENDplex kit (Biolegend) according to the manufacturer’s instructions and normalized by total protein content. Total protein content was quantified using the Pierce BCA Protein Assay kit (Thermo Fisher Scientific).

### Histological analysis of immune infiltrates in tumor, lung, liver and kidney

RM9-hSTEAP1-fluc tumor-bearing B6 mice received 5 million CAR-T cells on day 4 since tumor inoculation. Tumors, lungs, livers, and kidneys were collected on day 14 and fixed with 4% paraformaldehyde solution in PBS overnight at 4°C. FFPE tissue samples were sectioned followed by hematoxylin and eosin (H&E) staining and immunohistochemistry analysis as previously described^24^. Briefly, tissue slides were deparaffinized and rehydrated followed by antigen retrieval in Citrate-Based Antigen Unmasking Solution (Vector Labs). Slides were blocked and stained with rabbit anti-CD3 antibody (Thermo Fisher, MA5-14524, 1:100) for 1 h at 37 °C. Slides were washed three times with TBST (pH=8.0) and incubated with PowerVision Poly-HRP anti-rabbit IgG (Leica Biosystems) at 37 °C for 30 min. Post three washes slides were incubated with 3,3′-Diaminobenzidine (DAB) (Sigma Aldrich) at room temperature for 10 min. Slides were stained for H&E followed by dehydration steps and mounting. Stained slides were scanned at Translational Pathology Core Laboratory (TPCL) at UCLA. QuPath 0.2.3 was used to analyze the IHC images^44^.

### Analysis of immune cells in tumor using flow cytometry

RM9-hSTEAP1-fluc tumor-bearing mice received 5 million CAR-T cells on day 4 since tumor inoculation. Tumors were collected on day 14 and cut into small pieces using surgical scissors and digested in DMEM supplemented with 2% FBS, 2 mg/mL collagenase D (Sigma-Aldrich) and 40 µg/mL DNase I (Roche) for 30 min at 37°C. Single-cell suspensions in DMEM supplemented with 2%FBS were prepared from digested tumors using a 70 µm cell strainer (Thermo Fisher Scientific). Red blood cells were lysed with ACK lysing buffer (Gibco) for 5 minutes at room temperature and neutralized with PBS (-). Cells were stained with BD Horizon Fixable Viability Stain 510 (BD). Fc receptors were blocked using purified anti-mouse CD16/32 antibody (93, Biolegend). Cells were stained with a cocktail of anti-mouse antibodies and fixed with eBioscience IC Fixation Buffer (Invitrogen). Cells were acquired using a BD FACSymphony A3 flow cytometer and data were analyzed using FlowJo (BD). Simple linear regression was performed to analyze correlation between tumor burden and immune cell infiltrates using GraphPad Prism 10. The following anti-mouse antibodies were used: CD45.2 APC-Cy7 (30-F11, Biolegend), CD3 BUV395 (145-2C11, BD), CD4 BUV805 (GK1.5, BD), CD8 Alexa Fluor 700 (53-6.7, Biolegend), NK1.1 PerCP-Cy5.5 (PK136, Biolegend), PD-1 BV605 (29F.1A12, Biolegend), CTLA-4 APC (UC10-4B9, Biolegend), CD11b APC-Cy7 (M1/70, Invitrogen), Ly6G BUV737 (1A8, BD), Ly6C Alexa Fluor 488 (HK1.4, Biolegend), CD19 BV785 (6D5, Biolegend), CD11c PE-Cy7 (HL3, BD), MHC-II (I-A/I-E) BV711 (M5/114.15.2, Biolegend), F4/80 PE-Cy5 (BM8, Biolegend) and CD103 BV605 (2E7, Biolegend). Gating strategies are shown in Fig. S10 and S11.

### GeoMX High-plex transcriptome analysis

NanoString GeoMx Digital Spatial Profiler (DSP) whole transcriptome analysis (WTA) was performed on tumor sample collected at day 14 after tumor inoculation from hSTEAP1-mBBζ CAR T cell and hSTEAP1-mBBζ CAR +NFAT-CBD-IL-12 T cell treated groups. Parafin embedded tissue blocks were sectioned by UCLA Translational Pathology Core Laboratory (TPCL). NanoString GeoMx DSP was performed at Technology Center for Genomics & Bioinformatics (TCGB) core at UCLA. Twelve regions of interest (ROI) were chosen across four different tumor sections in each treated group followed by library preparation and sequencing. A tumor sample in unarmored CAR-T treatment group was not used because the tissue section was damaged.

Transcriptome data analysis was performed using NanoString Spatial Data Analysis software (GeoMx® DSP Software Version 3.0.0.109). Raw files were analyzed for quality control, followed by sequence alignment. Count matrix files were used to perform differential gene expression analysis using DESeq2^45^. For PCA analysis, Fragments Per Kilobase of transcript per Million mapped reads (FPKM) values were normalized by log2 + 1 transformation and PCA was plotted based on a correlation matrix using the prcomp package v3.6.2. PCA plots were visualized using the factoextra package v1.0.7 and ggpubr package v0.6.0. S1-12 was excluded as an outlier in GSEA based on the PCA analysis. Pathway analysis was performed using GSEA^46^. All computational analyses were carried out in RStudio v4.1.0. Heatmaps were generated using the package pheatmap v1.0.12. One-sided Fisher’s exact test was performed using fisher.test() function in R v4.2.1.

### Splenic T cell response to hSTEAP1 positive and negative RM9 cells after CAR-T treatment

RM9-hSTEAP1-fluc tumor-bearing mice received 10 million CAR-T cells on day 4 since tumor inoculation. Spleens were harvested on day 12 and splenic T cells were purified using MagniSort Mouse T cell Enrichment Kit (Invitrogen). The T cell enrichment cocktail was supplemented with 1 µg/mL Biotin-SP (long spacer) AffiniPure F(ab’)_2_ Fragment Donkey Anti-Human IgG (H+L) (Jackson ImmunoResearch, 709-066-149) to remove CAR-T cells. Isolated splenic T cells were maintained in the mouse T cell culture medium until use. A day before co-culture with splenic T cells, 3 × 10^6^ tumor cells were seeded in a T75 flask and supplemented with 20 ng/mL recombinant mouse IFN-γ (Biolegend) and incubated for 24 h at 37°C in the presence of 5% CO_2_. On the next day, tumor cells were harvested using TryPLE express Emzyme (Gibco) and washed twice with the mouse T cell culture medium without cytokines. 2 × 10^5^ tumor cells and the same number of splenic T cells were mixed in 200 µL of mouse T cell culture medium without cytokines and seeded in a 96-well V-bottom plate (Corning) and incubated for 48 h at 37°C in the presence of 5% CO_2_. After centrifuge (300 g, 5 min), Supernatant was collected and IFN-γ was quantified by mouse IFN-γ ELISA kit (Invitrogen).

### Manufacturing and functional testing of CAR Jurkat cells

Jurkat cells were transduced with 10x concentrated lentiviral vector using a Retronectin-coated plate as described above. Surface expression of the CAR was detected using Biotin-SP-conjugated AffiniPure F(ab’)2 Fragment Goat Anti-Human IgG, F(ab’)2 Fragment Specific (Jackson ImmunoResearch, 109-066-006) and PE Streptavidin (Biolegend). 50000 CAR^+^ Jurkat T cells were stimulated with eBioscience Cell Stimulation Cocktail (Invitrogen). Secreted IL-12 variants were quantified by Human IL-12 p70 DuoSet ELISA (R&D Systems). Human scIL-12-His and CBD-human scIL-12-His were used as standards.

### Manufacturing and functional testing of human CAR-T cells

Healthy donor human PBMC (STEMCELL Technologies) were subjected to EasySep CD8 negative selection (STEMCELL Technologies). 1 × 10^6^/mL CD8^+^ T cells were stimulated with T cell TransAct (1:100, Miltenyi Biotec) in RPMI media supplemented with 5% human serum (Bloodworks Northwest) and 50 U/mL human IL-2 (Peprotech). Next day, 5x concentrated lentiviral vector was added to the culture with 10 µg/mL protamine sulfate (MP Biomedicals). 24 hours later, media were replaced with fresh IL-2 to remove viruses and TransAct. Seven days after the addition of viruses, cells were stained with 1:100 biotin-anti-Fab (Jackson Immunoreseaerch, 109-066-006) followed by 1:200 PE-streptavidin (ThermoFisher Scientific), and CAR^+^ cells were sorted (except untransduced cells from which all live cells were sorted) on BD FACSymphony S6. 5 × 10^4^ sorted cells were expanded with irradiated feeder cells (2 × 10^6^ mixed PBMCs (three individual donor PBMCs from STEMCELL Technologies) and 4 × 10^5^ Epstein-Barr virus-transformed B-lymphoblastoid cells (Fred Hutch Research Cell Bank)), 50 U/mL IL-2, 10 ng/mL IL-15 (Peprotech), and 0.3 mL anti-CD3 antibodies (clone OKT3, Miltenyi Biotec) in 1 mL culture volume. Culture media (10% human serum and 50 U/mL IL-2-supplemented RPMI) were replaced every two or three days. After 10 days, T cells were used for functional assays. For the killing assay, 1 × 10^5^ T cells were co-cultured with 1 × 10^4^ target cells (22Rv1 hSTEAP1 KO + rescue or 22Rv1 hSTEAP1 KO)^24^ lentivirally transduced to express mCherry in 96-well flat-bottom plates (Corning). Target killing was monitored in IncuCyte (Sartorius), detecting fluorescence from mCherry every four hours using whole-well scanning. Integrated intensity per well was normalized by values from the first time point. For cytokine production, 1 × 10^5^ each T cells and target cells were co-cultured in 96-well round-bottom plates (Corning) with 1:1000-diluted Golgi Stop (monensin, BD) and Golgi Plug (brefeldin A, BD) for 18 hours. Cells were then surface-stained with Live/Dead Aqua (1:1000, ThermoFisher Scientific) and 1:200 FITC-CD8 (Clone SK1, 1:200, Biolegend) for 15 minutes at 4 °C, fixed and permeabilized with BD Cytofix/Cytoperm (BD), intracellular-stained with APC-IFN-γ (Clone 4S.B3, 1:100, Biolegend) and PE-IL-12 (Clone 20C2, 1:100, BD) for 15 minutes at 4 °C, and analyzed on BD FACSymphony A5.

### Statistical analysis

Statistical analysis was performed using GraphPad Prism 10. Unpaired t test with Welch’s correction (parametric data) or Mann-Whitney U test (non-parametric data determined based on F test) was used for comparisons between two groups. One-way analysis of variance (ANOVA) followed by Tukey’s multiple comparisons test (parametric data) or Kruskal-Wallis test followed by Dunn’s multiple comparison test (non-parametric data determined based on Brown-Forsythe test) was used for comparisons of groups of three or more except for the splenic T cell restimulation experiment where two-way ANOVA followed by Šídák’s multiple comparisons test was used. Log-rank (Mantel-Cox) test was used for comparisons of survival curves between two groups.

## Reporting Summary

Further information on research design is available in the Nature Research Reporting Summary linked to this article.

## Data availability

The main data supporting the results in this study are available within the paper and its Supplementary Information. Source data for the figures will be provided with this paper.

## Acknowledgements

We thank Deborah E. Banker for her helpful comments and editing. We thank the LMS/NIHR Imperial Biomedical Research Centre Flow Cytometry Facility for its support. This research was funded in part by JSPS Overseas Research Fellowships (202160429 to K.S.), European Molecular Biology Organization Postdoctoral Fellowships (ALTF 56-2022 to K.S.), Prostate Cancer Foundation Young Investigator Awards (to V.B.), Department of Defense Prostate Cancer Research Program Early Investigator Research Award (HT9425-23-1-0089 to V.B.), the Department of Defense Prostate Cancer Research Program Awards (W81XWH-21-1-0581 to J.K.L and J.I.), the Pacific Northwest Prostate Cancer SPORE P50 097186 (to J.K.L.), The Academy of Medical Sciences Springboard (SBF007\100097 to J.I.) and Prostate Cancer UK Research Innovation Awards (RIA21-ST2-010 to J.K.L. and J.I.).

## Author contributions

K.S., J.K.L and J.I. conceived the project. K.S., V.B., Y.A., S.J.P., A.G.C., J.K.L and J.I. designed experiments. K.S., V.B., Y.A. and J.B. performed experiments. P.K., C.W., M.N., T.M., O.D., P.-C.C. and T.C. assisted with tumor experiments. V.B. and G.J. analyzed spatial genomics data. K.S. and J.I. wrote the original draft. K.S., V.B., Y.A., S.J.P., A.G.C., J.K.L. and J.I. edited the manuscript with input from all other authors. J.K.L. and J.I. supervised the research.

## Additional Information

Requests for materials should be addressed to John K. Lee or Jun Ishihara.

