## Supplemental Table for "*Collagen-binding IL-12 expressing* STEAP1 CAR-T cells reduce toxicity and eradicate mouse prostate cancer in combination with checkpoint inhibitors"

**Affiliations:**

**Table S1. Amino acid sequences of the scIL-12 variants**

Mouse IL-12 (for production in HEK293F):

MWELEKDVYVVEVDWTPDAPGETVNLTCDTPEEDDITWTSDQRHGVIGSGKTLTITVKEFLDAGQYTCHKGGETLSHSHLLLHKKENGIWSTEILKNFKNKTFLKCEAPNYSGRFTCSWLVQRNMDLKFNIKSSSSSPDSRAVTCGMASLSAEKVTLDQRDYEKYSVSCQEDVTCPTAEETLPIELALEARQQNKYENYSTSFFIRDIIKPDPPKNLQMKPLKNSQVEVSWEYPDSWSTPHSYFSLKFFVRIQRKKEKMKETEEGCNQKGAFLVEKTSTEVQCKGGNVCVQAQDRYYNSSCSKWACVPCRVRSGGSGGGSGGGSGGGSRVIPVSGPARCLSQSRNLLKTTDDMVKTAREKLKHYSCTAEDIDHEDITRDQTSTLKTCLPLELHKNESCLATRETSSTTRGSCLPPQKTSLMMTLCLGSIYEDLKMYQTEFQAINAALQNHNHQQIILDKGMLVAIDELMQSLNHNGETLRQKPPVGEADPYRVKMKLCILLHAFSTRVVTINRVMGYLSSAHHHHHH*

Mouse IL-12 (as a payload for CAR-T):

MWELEKDVYVVEVDWTPDAPGETVNLTCDTPEEDDITWTSDQRHGVIGSGKTLTITVKEFLDAGQYTCHKGGETLSHSHLLLHKKENGIWSTEILKNFKNKTFLKCEAPNYSGRFTCSWLVQRNMDLKFNIKSSSSSPDSRAVTCGMASLSAEKVTLDQRDYEKYSVSCQEDVTCPTAEETLPIELALEARQQNKYENYSTSFFIRDIIKPDPPKNLQMKPLKNSQVEVSWEYPDSWSTPHSYFSLKFFVRIQRKKEKMKETEEGCNQKGAFLVEKTSTEVQCKGGNVCVQAQDRYYNSSCSKWACVPCRVRSGGSGGGSGGGSGGGSRVIPVSGPARCLSQSRNLLKTTDDMVKTAREKLKHYSCTAEDIDHEDITRDQTSTLKTCLPLELHKNESCLATRETSSTTRGSCLPPQKTSLMMTLCLGSIYEDLKMYQTEFQAINAALQNHNHQQIILDKGMLVAIDELMQSLNHNGETLRQKPPVGEADPYRVKMKLCILLHAFSTRVVTINRVMGYLSSA*

CBD-mouse IL-12 (for production in HEK293F):

CSQPLDVILLLDGSSSFPASYFDEMKSFAKAFISKANIGPRLTQVSVLQYGSITTIDVPWNVVPEKAHLLSLVDVMQREGGPSQIGDALGFAVRYLTSEMHGARPGASKAVVILVTDVSVDSVDAAADAARSNRVTVFPIGIGDRYDAAQLRILAGPAGDSNVVKLQRIEDLPTMVTLGNSFLHKLCSGFVRIGGGSGGGSMWELEKDVYVVEVDWTPDAPGETVNLTCDTPEEDDITWTSDQRHGVIGSGKTLTITVKEFLDAGQYTCHKGGETLSHSHLLLHKKENGIWSTEILKNFKNKTFLKCEAPNYSGRFTCSWLVQRNMDLKFNIKSSSSSPDSRAVTCGMASLSAEKVTLDQRDYEKYSVSCQEDVTCPTAEETLPIELALEARQQNKYENYSTSFFIRDIIKPDPPKNLQMKPLKNSQVEVSWEYPDSWSTPHSYFSLKFFVRIQRKKEKMKETEEGCNQKGAFLVEKTSTEVQCKGGNVCVQAQDRYYNSSCSKWACVPCRVRSGGSGGGSGGGSGGGSRVIPVSGPARCLSQSRNLLKTTDDMVKTAREKLKHYSCTAEDIDHEDITRDQTSTLKTCLPLELHKNESCLATRETSSTTRGSCLPPQKTSLMMTLCLGSIYEDLKMYQTEFQAINAALQNHNHQQIILDKGMLVAIDELMQSLNHNGETLRQKPPVGEADPYRVKMKLCILLHAFSTRVVTINRVMGYLSSAHHHHHH*

CBD-mouse IL-12 (as a payload for CAR-T):

CSQPLDVILLLDGSSSFPASYFDEMKSFAKAFISKANIGPRLTQVSVLQYGSITTIDVPWNVVPEKAHLLSLVDVMQREGGPSQIGDALGFAVRYLTSEMHGARPGASKAVVILVTDVSVDSVDAAADAARSNRVTVFPIGIGDRYDAAQLRILAGPAGDSNVVKLQRIEDLPTMVTLGNSFLHKLCSGFVRIGGGSGGGSMWELEKDVYVVEVDWTPDAPGETVNLTCDTPEEDDITWTSDQRHGVIGSGKTLTITVKEFLDAGQYTCHKGGETLSHSHLLLHKKENGIWSTEILKNFKNKTFLKCEAPNYSGRFTCSWLVQRNMDLKFNIKSSSSSPDSRAVTCGMASLSAEKVTLDQRDYEKYSVSCQEDVTCPTAEETLPIELALEARQQNKYENYSTSFFIRDIIKPDPPKNLQMKPLKNSQVEVSWEYPDSWSTPHSYFSLKFFVRIQRKKEKMKETEEGCNQKGAFLVEKTSTEVQCKGGNVCVQAQDRYYNSSCSKWA

CVPCRVRSGGSGGGSGGGSGGGSRVIPVSGPARCLSQSRNLLKTTDDMVKTAREKLKHYSCTAEDIDHEDITRDQTSTLKTCLPLELHKNESCLATRETSSTTRGSCLPPQKTSLMMTLCLGSIYEDLKMYQTEFQAINAALQNHNHQQIILDKGMLVAIDELMQSLNHNGETLRQKPPVGEADPYRVKMKLCILLHAFSTRVVTINRVMGYLSSA*

Mouse IL-12-CBD (for production in HEK293F):

MWELEKDVYVVEVDWTPDAPGETVNLTCDTPEEDDITWTSDQRHGVIGSGKTLTITVKEFLDAGQYTCHKGGETLSHSHLLLHKKENGIWSTEILKNFKNKTFLKCEAPNYSGRFTCSWLVQRNMDLKFNIKSSSSSPDSRAVTCGMASLSAEKVTLDQRDYEKYSVSCQEDVTCPTAEETLPIELALEARQQNKYENYSTSFFIRDIIKPDPPKNLQMKPLKNSQVEVSWEYPDSWSTPHSYFSLKFFVRIQRKKEKMKETEEGCNQKGAFLVEKTSTEVQCKGGNVCVQAQDRYYNSSCSKWACVPCRVRSGGSGGGSGGGSGGGSRVIPVSGPARCLSQSRNLLKTTDDMVKTAREKLKHYSCTAEDIDHEDITRDQTSTLKTCLPLELHKNESCLATRETSSTTRGSCLPPQKTSLMMTLCLGSIYEDLKMYQTEFQAINAALQNHNHQQIILDKGMLVAIDELMQSLNHNGETLRQKPPVGEADPYRVKMKLCILLHAFSTRVVTINRVMGYLSSACSQPLDVILLLDGSSSFPASYFDEMKSFAKAFISKANIGPRLTQVSVLQYGSITTIDVPWNVVPEKAHLLSLVDVMQREGGPSQIGDALGFAVRYLTSEMHGARPGASKAVVILVTDVSVDSVDAAADAARSNRVTVFPIGIGDRYDAAQLRILAGPAGDSNVVKLQRIEDLPTMVTLGNSFLHKLCSGFVRIGGGSGGGSHHHHHH*

Mouse IL-12-CBD (as a payload for CAR-T):

MWELEKDVYVVEVDWTPDAPGETVNLTCDTPEEDDITWTSDQRHGVIGSGKTLTITVKEFLDAGQYTCHKGGETLSHSHLLLHKKENGIWSTEILKNFKNKTFLKCEAPNYSGRFTCSWLVQRNMDLKFNIKSSSSSPDSRAVTCGMASLSAEKVTLDQRDYEKYSVSCQEDVTCPTAEETLPIELALEARQQNKYENYSTSFFIRDIIKPDPPKNLQMKPLKNS

QVEVSWEYPDSWSTPHSYFSLKFFVRIQRKKEKMKETEEGCNQKGAFLVEKTSTEVQCKGGNVCVQAQDRYYNSSCSKWACVPCRVRSGGSGGGSGGGSGGGSRVIPVSGPARCLSQSRNLLKTTDDMVKTAREKLKHYSCTAEDIDHEDITRDQTSTLKTCLPLELHKNESCLATRETSSTTRGSCLPPQKTSLMMTLCLGSIYEDLKMYQTEFQAINAALQNHNHQQIILDKGMLVAIDELMQSLNHNGETLRQKPPVGEADPYRVKMKLCILLHAFSTRVVTINRVMGYLSSACSQPLDVILLLDGSSSFPASYFDEMKSFAKAFISKANIGPRLTQVSVLQYGSITTIDVPWNVVPEKAHLLSLVDVMQREGGPSQIGDALGFAVRYLTSEMHGARPGASKAVVILVTDVSVDSVDAAADAARSNRVTVFPIGIGDRYDAAQLRILAGPAGDSNVVKLQRIEDLPTMVTLGNSFLHKLCSGFVRI*

CBD-mouse IL-12-CBD (for production in HEK293F):

CSQPLDVILLLDGSSSFPASYFDEMKSFAKAFISKANIGPRLTQVSVLQYGSITTIDVPWNVVPEKAHLLSLVDVMQREGGPSQIGDALGFAVRYLTSEMHGARPGASKAVVILVTDVSVDSVDAAADAARSNRVTVFPIGIGDRYDAAQLRILAGPAGDSNVVKLQRIEDLPTMVTLGNSFLHKLCSGFVRIGGGSGGGSMWELEKDVYVVEVDWTPDAPGETVNLTCDTPEEDDITWTSDQRHGVIGSGKTLTITVKEFLDAGQYTCHKGGETLSHSHLLLHKKENGIWSTEILKNFKNKTFLKCEAPNYSGRFTCSWLVQRNMDLKFNIKSSSSSPDSRAVTCGMASLSAEKVTLDQRDYEKYSVSCQEDVTCPTAEETLPIELALEARQQNKYENYSTSFFIRDIIKPDPPKNLQMKPLKNSQVEVSWEYPDSWSTPHSYFSLKFFVRIQRKKEKMKETEEGCNQKGAFLVEKTSTEVQCKGGNVCVQAQDRYYNSSCSKWACVPCRVRSGGSGGGSGGGSGGGSRVIPVSGPARCLSQSRNLLKTTDDMVKTAREKLKHYSCTAEDIDHEDITRDQTSTLKTCLPLELHKNESCLATRETSSTTRGSCLPPQKTSLMMTLCLGSIYEDLKMYQTEFQAINAALQNHNHQQIILDKGMLVAIDELMQSLNHNGETLRQKPPVGEADPYRVKMKLCILLHAFSTRVVTINRVMGYLSSACSQPLDVILLLDGSSSFPASYFDEMKSFAKAFISKANIGPRLTQVSVLQYGSITTIDVPWNVVPEKA

HLLSLVDVMQREGGPSQIGDALGFAVRYLTSEMHGARPGASKAVVILVTDVSVDSVDAAADAARSNRVTVFPIGIGDRYDAAQLRILAGPAGDSNVVKLQRIEDLPTMVTLGNSFLHKLCSGFVRIGGGSGGGSHHHHHH*

CBD-mouse IL-12-CBD (as a payload for CAR-T):

CSQPLDVILLLDGSSSFPASYFDEMKSFAKAFISKANIGPRLTQVSVLQYGSITTIDVPWNVVPEKAHLLSLVDVMQREGGPSQIGDALGFAVRYLTSEMHGARPGASKAVVILVTDVSVDSVDAAADAARSNRVTVFPIGIGDRYDAAQLRILAGPAGDSNVVKLQRIEDLPTMVTLGNSFLHKLCSGFVRIGGGSGGGSMWELEKDVYVVEVDWTPDAPGETVNLTCDTPEEDDITWTSDQRHGVIGSGKTLTITVKEFLDAGQYTCHKGGETLSHSHLLLHKKENGIWSTEILKNFKNKTFLKCEAPNYSGRFTCSWLVQRNMDLKFNIKSSSSSPDSRAVTCGMASLSAEKVTLDQRDYEKYSVSCQEDVTCPTAEETLPIELALEARQQNKYENYSTSFFIRDIIKPDPPKNLQMKPLKNSQVEVSWEYPDSWSTPHSYFSLKFFVRIQRKKEKMKETEEGCNQKGAFLVEKTSTEVQCKGGNVCVQAQDRYYNSSCSKWACVPCRVRSGGSGGGSGGGSGGGSRVIPVSGPARCLSQSRNLLKTTDDMVKTAREKLKHYSCTAEDIDHEDITRDQTSTLKTCLPLELHKNESCLATRETSSTTRGSCLPPQKTSLMMTLCLGSIYEDLKMYQTEFQAINAALQNHNHQQIILDKGMLVAIDELMQSLNHNGETLRQKPPVGEADPYRVKMKLCILLHAFSTRVVTINRVMGYLSSACSQPLDVILLLDGSSSFPASYFDEMKSFAKAFISKANIGPRLTQVSVLQYGSITTIDVPWNVVPEKAHLLSLVDVMQREGGPSQIGDALGFAVRYLTSEMHGARPGASKAVVILVTDVSVDSVDAAADAARSNRVTVFPIGIGDRYDAAQLRILAGPAGDSNVVKLQRIEDLPTMVTLGNSFLHKLCSGFVRI*

Human IL-12 (for production in HEK293F):

IWELKKDVYVVELDWYPDAPGEMVVLTCDTPEEDGITWTLDQSSEVLGSGKTLTIQVKEFGDAGQYTCHKGGEVLSHSLLLLHKKEDGIWSTDILKDQKEPKNKTFLRCEAKNYSGRFTCWWLTTISTDLTFSVKSSRGSSDPQGVTCGAATLSAERVRGDNKEYEYSVECQEDSACPAAEESLPIEVMVDAVHKLKYENYTSSFFIRDIIKPDPPKNLQLKPLKNSRQVEVSWEYPDTWSTPHSYFSLTFCVQVQGKSKREKKDRVFTDKTSATVICRKNASISVRAQDRYYSSSWSEWASVPCSGGGGSGGGGSGGGGSRNLPVATPDPGMFPCLHHSQNLLRAVSNMLQKARQTLEFYPCTSEEIDHEDITKDKTSTVEACLPLELTKNESCLNSRETSFITNGSCLASRKTSFMMALCLSSIYEDLKMYQVEFKTMNAKLLMDPKRQIFLDQNMLAVIDELMQALNFNSETVPQKSSLEEPDFYKTKIKLCILLHAFRIRAVTIDRVMSYLNASHHHHHH*

Human IL-12 (as a payload for CAR-T):

IWELKKDVYVVELDWYPDAPGEMVVLTCDTPEEDGITWTLDQSSEVLGSGKTLTIQVKEFGDAGQYTCHKGGEVLSHSLLLLHKKEDGIWSTDILKDQKEPKNKTFLRCEAKNYSGRFTCWWLTTISTDLTFSVKSSRGSSDPQGVTCGAATLSAERVRGDNKEYEYSVECQEDSACPAAEESLPIEVMVDAVHKLKYENYTSSFFIRDIIKPDPPKNLQLKPLKNSRQVEVSWEYPDTWSTPHSYFSLTFCVQVQGKSKREKKDRVFTDKTSATVICRKNASISVRAQDRYYSSSWSEWASVPCSGGGGSGGGGSGGGGSRNLPVATPDPGMFPCLHHSQNLLRAVSNMLQKARQTLEFYPCTSEEIDHEDITKDKTSTVEACLPLELTKNESCLNSRETSFITNGSCLASRKTSFMMALCLSSIYEDLKMYQVEFKTMNAKLLMDPKRQIFLDQNMLAVIDELMQALNFNSETVPQKSSLEEPDFYKTKIKLCILLHAFRIRAVTIDRVMSYLNAS*

CBD-human IL-12 (for production in HEK293F):

CSQPLDVILLLDGSSSFPASYFDEMKSFAKAFISKANIGPRLTQVSVLQYGSITTIDVPWNVVPEKAHLLSLVDVMQREGGPSQIGDALGFAVRYLTSEMHGARPGASKAVVILVTDVSVDSVDAAADAARSNRVTVFPIGIGDRYDAAQLRILAGPAGDSNVVKLQRIEDLPTMVTLGNSFLHKLCSGFVRIGGGSGGGSIWELKKDVYVVELDWYPDAPGEMVVLTCDTPEEDGITWTLDQSSEVLGSGKTLTIQVKEFGDAGQYTCHKGGEVLSHSLLLLHKKEDGIWSTDILKDQKEPKNKTFLRCEAKNYSGRFTCWWLTTISTDLTFSVKSSRGSSDPQGVTCGAATLSAERVRGDNKEYEYSVECQEDSACPAAEESLPIEVMVDAVHKLKYENYTSSFFIRDIIKPDPPKNLQLKPLKNSRQVEVSWEYPDTWSTPHSYFSLTFCVQVQGKSKREKKDRVFTDKTSATVICRKNASISVRAQDRYYSSSWSEWASVPCSGGGGSGGGGSGGGGSRNLPVATPDPGMFPCLHHSQNLLRAVSNMLQKARQTLEFYPCTSEEIDHEDITKDKTSTVEACLPLELTKNESCLNSRETSFITNGSCLASRKTSFMMALCLSSIYEDLKMYQVEFKTMNAKLLMDPKRQIFLDQNMLAVIDELMQALNFNSETVPQKSSLEEPDFYKTKIKLCILLHAFRIRAVTIDRVMSYLNASHHHHHH*

CBD-human IL-12 (as a payload for CAR-T):

CSQPLDVILLLDGSSSFPASYFDEMKSFAKAFISKANIGPRLTQVSVLQYGSITTIDVPWNVVPEKAHLLSLVDVMQREGGPSQIGDALGFAVRYLTSEMHGARPGASKAVVILVTDVSVDSVDAAADAARSNRVTVFPIGIGDRYDAAQLRILAGPAGDSNVVKLQRIEDLPTMVTLGNSFLHKLCSGFVRIGGGSGGGSIWELKKDVYVVELDWYPDAPGEMVVLTCDTPEEDGITWTLDQSSEVLGSGKTLTIQVKEFGDAGQYTCHKGGEVLSHSLLLLHKKEDGIWSTDILKDQKEPKNKTFLRCEAKNYSGRFTCWWLTTISTDLTFSVKSSRGSSDPQGVTCGAATLSAERVRGDNKEYEYSVECQEDSACPAAEESLPIEVMVDAVHKLKYENYTSSFFIRDIIKPDPPKNLQLKPLKNSRQVEVSWEYPDTWSTPHSYFSLTFCVQVQGKSKREKKDRVFTDKTSATVICRKNASISVRAQDRYYSSSWSEWASVPCSGGGGSGGGGSGGGGSRNLPVATPDPGMFPCLHHSQNLLRAVSNMLQKARQTL

EFYPCTSEEIDHEDITKDKTSTVEACLPLELTKNESCLNSRETSFITNGSCLASRKTS
FMMALCLSSIYEDLKMYQVEFKTMNAKLLMDPKRQIFLDQNMLAVIDELMQALNFN
SETVPQKSSLEEPDFYKTKIKLCILLHAFRIRAVTIDRVMSYLNAS*
